# Capsid-specialized protein language models reveal higher-order viral architecture from sequence

**DOI:** 10.64898/2026.09.06.749605

**Authors:** Sihang Liu, Pingfeng Yu, Siqing Xia, Hong Wang

## Abstract

Viral capsid proteins preserve information on higher-order shell architecture and deep evolutionary history, yet current capsid annotation relies predominantly on homology-based methods that have reduced sensitivity across highly divergent environmental sequences. Here we develop ESMCapsid, a capsid-specialized protein language model for remote capsid detection and architecture-aware representation learning. Screening 343 million representative metagenomic protein clusters revealed a large homology-dark capsid repertoire, with approximately 62% of candidates lacking matches to existing reference databases. Sparse autoencoder decomposition identified recurrent semantic motifs linking homology-dark proteins to known structural lineages, suggesting that interpretable higher-order architectural information can be recovered directly from capsid sequences at metagenomic scale. Mapping conserved motif cores onto resolved viral shells showed spatial clustering and restricted radial positions, indicating that ESMCapsid captures geometric constraints beyond sequence similarity alone. Together, our findings establish a sequence-based route to organize homology-dark viral diversity through conserved architectural principles, extending viral discovery beyond sequence homology.

---

Viruses are abundant and ecologically important components of microbial ecosystems [1, 2], and their capsids constitute the principal structural shell of virions. Capsid proteins form the genome-enclosing structural shell of virions and serve as informative hallmarks for identifying viruses, assigning taxonomy, and characterizing viral resilience in the environment. [3, 4] Moreover, their structural folds can remain conserved even after primary-sequence similarity has become undetectable, making them especially useful for detecting remote evolutionary relationships among viruses. [5, 6] Structure-based studies have therefore helped define broad capsid lineages and connect viruses that are difficult to relate by sequence alone. Consequently, improving the annotation of viral capsid proteins is a necessary step toward characterizing the breadth of viral diversity and placing newly recovered viral sequences within meaningful structural and evolutionary contexts.

However, the diversity and characteristics of capsid proteins recovered from metagenomic datasets remain largely incomplete. [6–8] Current approaches still rely heavily on homology-based tools such as BLAST and profile hidden Markov models (HMMs), which lose sensitivity when close homologues are absent or query sequences are highly divergent. [9–11] Consequently, many divergent capsid proteins may remain unrecognized, leaving substantial regions of capsid-related sequence space unexplored. Although structure-based comparison can reveal remote relationships, exhaustive structure prediction and comparison remain computationally costly at the scale of modern metagenomic protein catalogues. [12, 13] This motivates scalable, interpretable and structure-aware representations for prioritizing and comparing candidate capsid proteins without modelling every sequence in three dimensions.

Protein language models (PLMs) offer one route to such representations because they map protein sequences to learned embeddings that encode aspects of structure and function. [12, 14–16] Recent PLM-based approaches have improved viral protein annotation and remote protein discovery [9, 11, 17], but their application to viral capsids raises two related challenges. First, general-purpose PLMs such as ESM and ProtTrans are trained on broad protein corpora in which viral capsid proteins constitute a small and highly specialized subset. [14, 15] It therefore remains unclear whether these models can faithfully represent the architectural diversity of remote capsids, rather than merely distinguish capsid proteins from non-capsid proteins. Second, most PLM-based analyses use dense, mean-pooled embeddings as global sequence descriptors. Although effective for classification, such embeddings do not directly expose the local and modular features that may encode fold conservation, lineage divergence and shell-level architectural constraints. [18, 19] Thus, a capsid-focused framework should combine sensitive remote recognition with a local, interpretable representation that remains comparable beyond detectable sequence homology.

To address this gap, we develop a capsid-specialized PLM framework that integrates remote capsid detection at metagenomic scale with semantic-motif-based representation learning. We first show that four general-purpose PLMs can effectively distinguish capsid proteins from non-capsid proteins, but do not fully capture their within-group diversity. We then derive two complementary ESMC-based models: ESMCapsid-S for sensitive remote capsid detection and ESMCapsid-C for capsid-specific representation learning. To interpret ESMCapsid-C representations, we decompose residue-level embeddings with sparse autoencoders and cluster token-level activation patterns into semantic motif tokens. Unlike conventional sequence motifs, these tokens are not defined by residue consensus. Instead, they capture recurrent activation contexts that remain comparable across proteins with little detectable sequence homology. Their n-gram profiles organize divergent capsids into broad capsid architecture categories and provide evidence for relationships between homology-dark groups and known structural lineages. Complementary structural analyses prioritize candidates with unresolved architectures and identify recurrent shell regions associated with capsid geometry. When applied to 343 million representative metagenomic protein clusters, this framework detects capsid candidates beyond the reach of profile-HMM searches and reveals habitat-structured reservoirs of capsid diversity that were previously poorly annotated. Together, this framework links metagenomic-scale remote capsid detection to architecture-informed interpretation through a local semantic-motif representation that remains comparable beyond detectable sequence homology.

## Results

### Generic PLMs support binary capsid recognition but incompletely represent within-capsid diversity

Previous studies have shown that PLMs are effective for diverse protein recognition and annotation. [12, 14, 20] To determine how well pretrained PLMs represent viral capsids, we benchmarked four models (three ESM series models and one ProFluent model) on capsid proteins, cellular proteins and challenging viral non-capsid proteins, including other structural proteins. Performance was evaluated under random, low-identity-held-out (<30% sequence identity) and family-held-out splits (Figure 1a and Figure 2a). Under five-fold cross-validation on the random split evaluation, all four models distinguished capsid proteins from non-capsid proteins with high accuracy (F1 > 0.94), and t-SNE projections [21] placed capsid proteins in a distinct region of embedding space (Extended Data Figure 1). However, capsid sequences consistently exhibited higher pseudoperplexity and masked-language-modelling (MLM) loss than non-capsid proteins, and classification performance declined more sharply on held-out families and low-identity samples (Figure 2c). Together, these results reveal a gap between accurate binary capsid/non-capsid discrimination and faithful representation of within-capsid diversity, indicating the need for capsid-specialized models for remote detection and representation learning.

**Figure 1.**
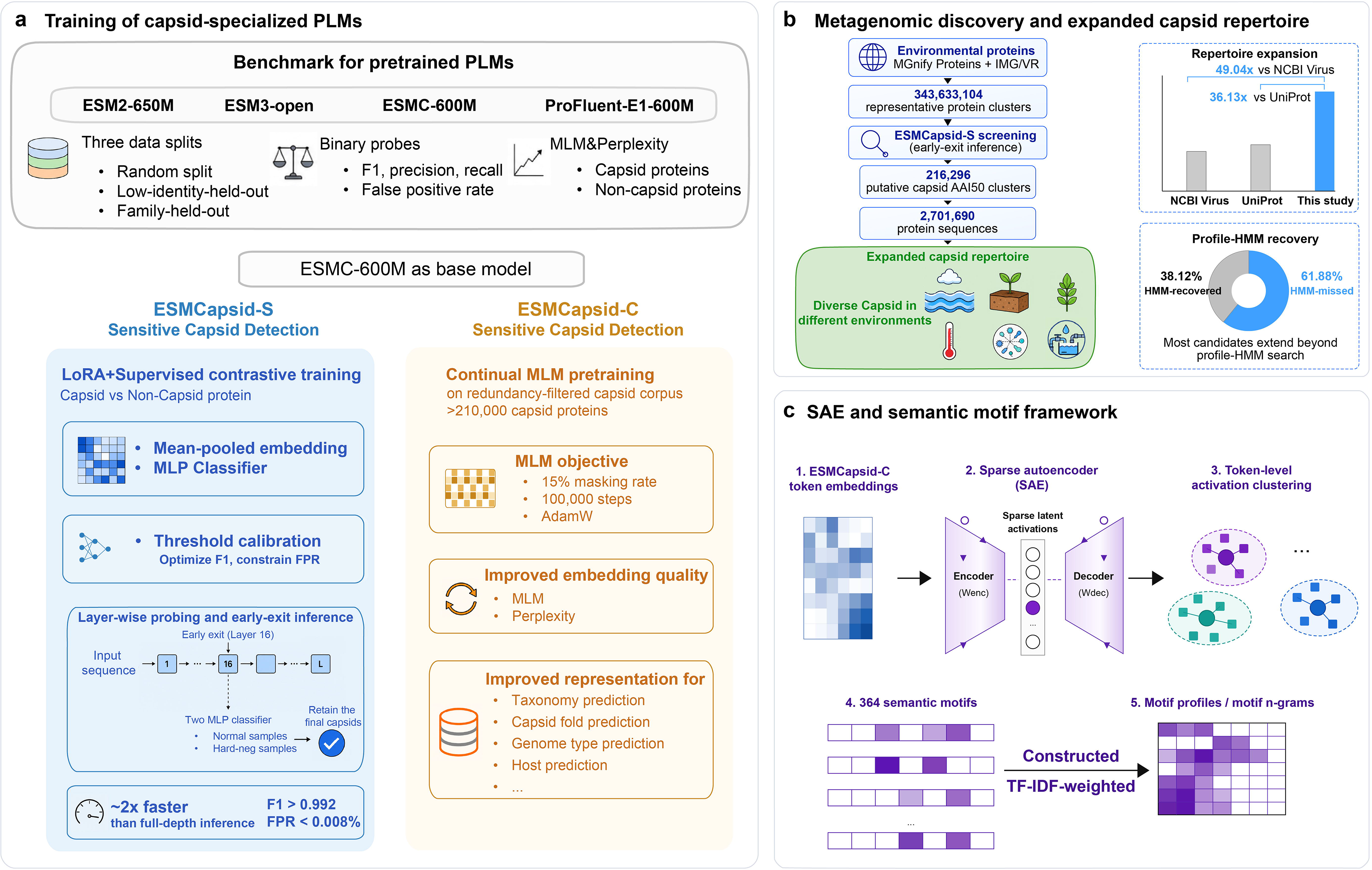
Development and application of capsid-specialized protein language models for metagenomic capsid discovery. (a) Workflow for training and evaluating capsid-specialized PLMs. (b) Large-scale metagenomic screening expands the detectable capsid repertoire. Environmental protein clusters derived from MGnify proteins and IMG/VR were screened with ESMCapsid-S, yielding 216,296 putative capsid AAI50 clusters comprising 2,701,690 protein sequences. (c) Sparse autoencoder-based interpretation of capsid embeddings. Token embeddings from the capsid-specialized model are decomposed into sparse latent activations using a sparse autoencoder, followed by token-level activation clustering to define semantic motifs. These motifs are summarized as TF-IDF-weighted motif profiles and motif n-grams, enabling interpretable analysis of capsid sequence features across the expanded metagenomic repertoire.

**Figure 2.**
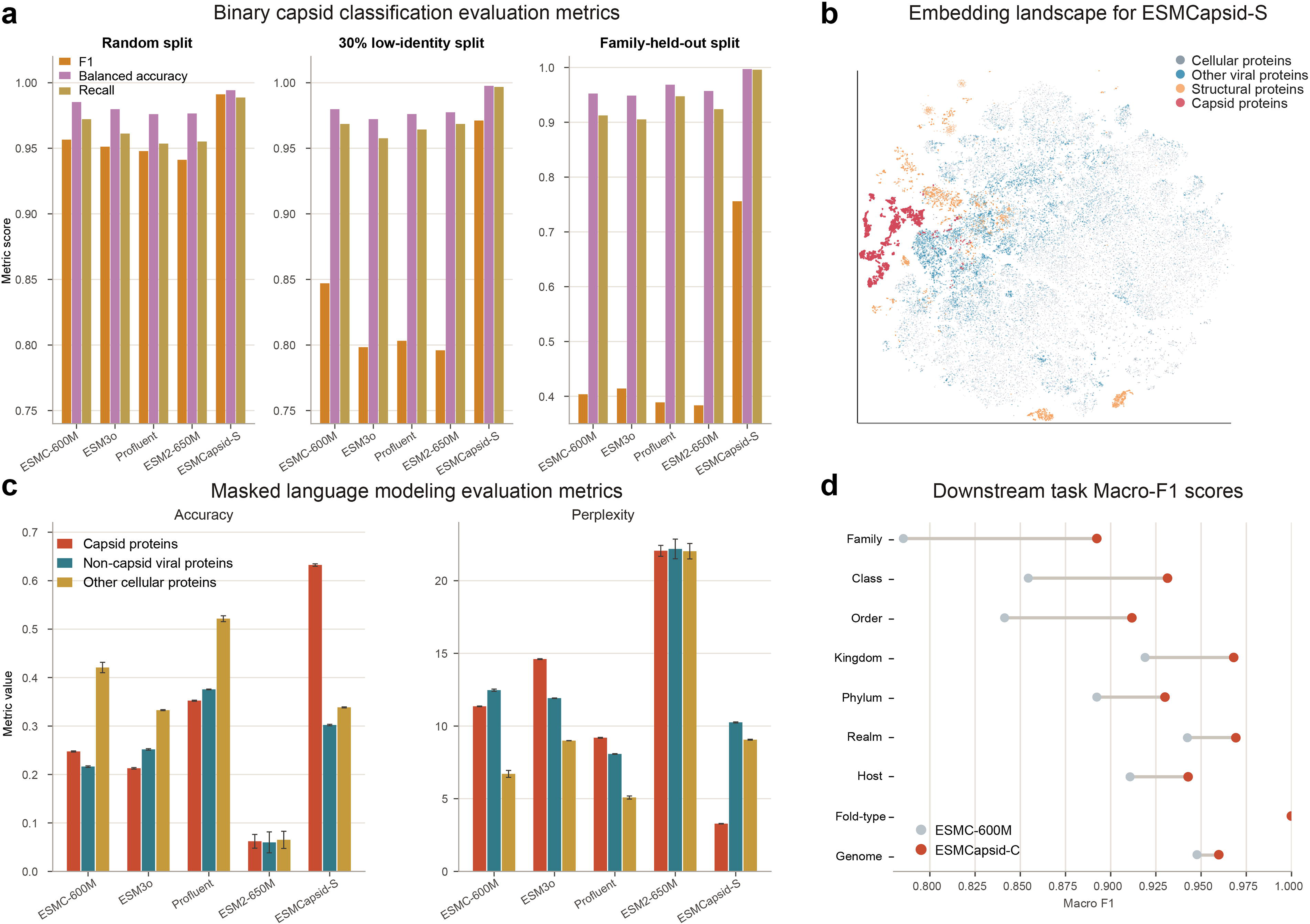
Benchmarking and representation properties of ESMCapsid models. (a) Binary capsid classification performance of four baseline protein language models and the tuned ESMCapsid-S model across three evaluation settings: random split, 30% low-identity holdout, and family-heldout generalization. For each model, bars show capsid F1 score, balanced accuracy, and recall. (b) Two-dimensional t-SNE projection of ESMCapsid-S embeddings, highlighting the distribution of capsid proteins relative to three background protein groups: other viral proteins, structural proteins, and cellular proteins. (c) Masked language modelling performance of four baseline models and ESMCapsid-C on three sequence groups: capsid proteins, non-capsid viral proteins, and cellular proteins. (d) Downstream task performance before and after ESMCapsid-C fine-tuning, shown as macro-F1 scores for taxonomic (Realm, Kingdom, Phylum, Class, Order, Family), host category (Animals, Bacteria, Plants, OtherEukaryote), capsid architecture category and genome-label classification tasks. Lines connect preand post-fine-tuning performance for each task.

### Capsid-specialized models improve remote detection and representation learning

We therefore selected the best-performing baseline model, ESMC, and fine-tuned it for binary capsid identification, yielding ESMCapsid-S (Figure 1a, see Methods). Layer-wise linear probing showed that capsid and non-capsid proteins were separable across layers (Extended Data Figure 2). Importantly, the gains of ESMCapsid-S were most pronounced under family-held-out and low-identity evaluation, where the model retained substantially higher recall, lower false-positive rate, and clearer separation in t-SNE space than the baseline model (Figure 2a and 2b), indicating improved out-of-distribution (OOD) detection of remote capsid proteins. To support large-scale screening, we implemented a Layer 16 dual-head screening strategy based on a shared intermediate representation. A primary classifier with a stringent capsid-probability threshold retained high-scoring capsid candidates, which were then re-evaluated by a classifier trained on hard negatives. This strategy maintained an F1 score above 0.992 and a false-positive rate of 0.008%, and reduced inference time by approximately two-fold relative to full-depth inference (Extended Data Figure 2).

Next, to improve representation quality for capsid proteins, we continued pretraining ESMC on a redundancy-filtered set of >210,000 capsid proteins compiled from UniProt, NCBI Virus, IMG/VR and MGnify Protein databases using a masked language modelling objective, yielding the capsid-specific model ESMCapsid-C (see Methods). Relative to the baseline model, ESMCapsid-C substantially reduced pseudo-perplexity and MLM loss on held-out capsid sequences, with only minor degradation on non-capsid proteins (Figure 2c), indicating effective domain adaptation without catastrophic forgetting. We then used layer-wise linear probes to assess the biological information encoded in the representations for four annotation tasks: capsid taxonomy, capsid architecture category, viral genome type and host category. ESMCapsid-C outperformed the baseline across the evaluated tasks, with particularly large gains at the family-level taxonomic classification (Figure 2d). Together, these results establish complementary roles for the two models: ESMCapsid-S provides high-throughput remote capsid detection, whereas ESMCapsid-C supports downstream capsid-specific representation learning and biological interpretation.

### Metagenomic-scale screening reveals an expanded and embedding-organized capsid repertoire

To systematically characterize capsid diversity from DNA and RNA viruses across diverse metagenomic samples represented in MGnify Proteins and IMG/VR, we first clustered more than 830 million protein sequences at 50% amino-acid identity (AAI50), yielding 343,633,104 AAI50 protein clusters. Each AAI50 representative was scored with the ESMCapsid-S dual-head classifier, which identified 216,296 putative capsid AAI50 clusters comprising 2,701,690 AAI90-dereplicated member sequences (Figure 3a). By sequence count, this repertoire was 49.04-fold larger than the capsid collection from NCBI Virus (55,089 sequences) and 36.13-fold larger than that from UniProt (74,780 sequences). Conventional profile-HMM searches recovered only 38.12% of these AAI50 clusters, and all HMM-recovered clusters were contained within our predictions (Figure 1b). Notably, only 74 protein clusters, corresponding to 0.03% of the predicted capsid AAI50 clusters, were flagged as potential non-capsid or cellular contaminants and excluded from downstream analyses. These results indicate that currently accessible environmental catalogues harbour a vast and still incompletely identified capsid repertoire that extends well beyond the reach of existing homology-based references.

**Figure 3.**
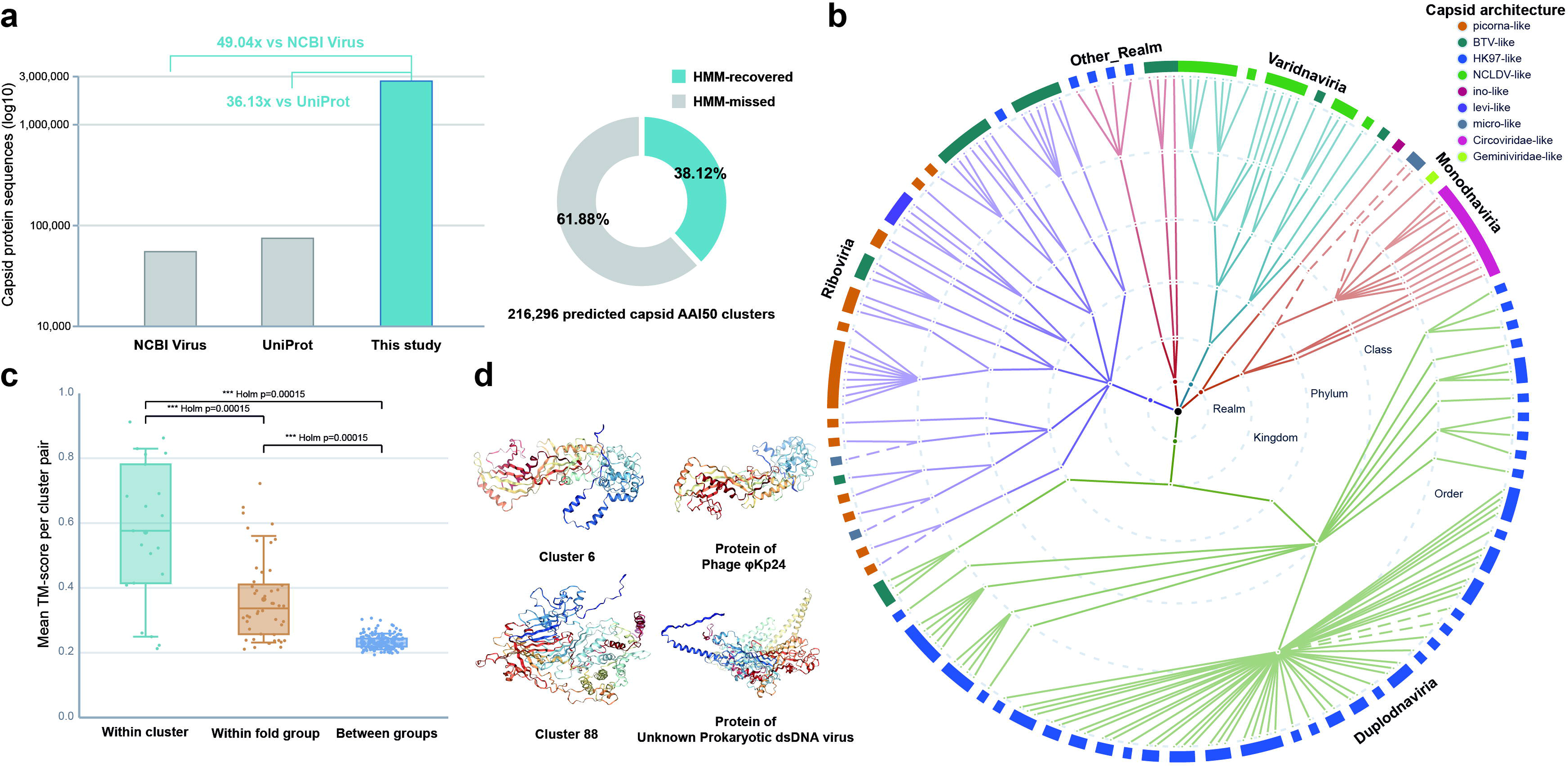
Capsid repertoire size, taxonomic distribution, structural similarity, and representative models. (a) Counts of capsid protein sequences from NCBI Virus, UniProt, and this study, shown on a log10 scale. (b) Radial taxonomic tree of capsid proteins across the viral taxonomic hierarchy, with the outer ring coloured by capsid architecture category. (c) Distribution of pairwise structural similarity, measured as the mean TM-score for cluster pairs within the same cluster, between clusters assigned to the same architecture category, and between architecture categories. Points represent individual cluster pairs, and Holm-adjusted pairwise significance is indicated. (d) Representative structural models for Cluster 6 and Cluster 88, together with their most structurally similar protein structures identified by Foldseek.

We next asked how this expanded capsid repertoire is organized within the capsid-specialized PLM latent space. Unsupervised clustering of pooled sequence-level embeddings generated by ESMCapsid-C resolved 99 embedding-defined capsid groups (see Methods), providing a latent-space map of the expanded capsid repertoire. To anchor this map within known viral diversity, we projected 13,504 reference major capsid proteins from NCBI Virus in the same embedding space (Figure 3b). Reference proteins mapped to 90 of the 99 groups, and the mean per-group dominant-taxon fractions were 97%, 97%, 94% and 93% at the realm, kingdom, phylum and class levels, respectively. By integrating embedding-based classification with HMM alignments, we confidently assigned 97 of the 99 embedding-defined groups to nine study-defined capsid architecture categories, including HK97-like (67 clusters), picorna-like (11), NCLDV-like (7), BTV-like (4), micro-like (3), Geminiviridae-like (2), ino-like (1), Circoviridae-like (1), and levi-like (1). This organization was not simply a recapitulation of existing profile-HMM annotations: Profile-HMM-supported architecture assignments were available for 77 of the 99 embedding-defined groups, leaving 22 clusters without direct HMM support. For 20 of the 22 clusters that can be assigned to a capsid architecture category by embedding, AlphaFold3 predictions for five randomly selected sequences per cluster showed that proteins within the same embedding-defined cluster were structurally coherent despite limited residue-level homology (Figure 3c). Within-group structural similarity (mean TM-score = 0.560) was significantly higher than similarity between embedding-defined groups assigned to the same architecture category (mean TM-score = 0.326) or to different architecture categories (mean TM-score = 0.256, Holm-adjusted *P <* 0.001 for all comparisons). Notably, two embedding-defined groups (groups 6 and 88) could not be assigned confidently to any capsid architecture category. BLAST and Foldseek searches [13, 22] indicated that group 6 is related to a recently described HK97-derived giant-phage capsid with a unique assembly strategy. [23, 24] Group 88 lacked a confident match to known capsid or non-capsid structures and therefore remains a candidate unresolved architecture among prokaryotic dsDNA viruses (Figure 3d). Together, these results show that the ESMCapsid-C embedding space organizes the expanded capsid repertoire into biologically coherent clusters, some of which are poorly resolved by conventional sequence-similarity and profile-HMM approaches, and has the potential to uncover previously unrecognized capsid fold diversity.

### Semantic motif profiles organize homology-dark capsids and link sequence representations to shell geometry

To interpret the embedding-defined groups, we decomposed residue-level ESMCapsid-C embeddings with a sparse autoencoder (SAE). [18, 19, 25] Leiden clustering of the resulting token-level activation vectors yielded 364 recurrent activation-pattern clusters, which we term semantic motif tokens (Figure 1c; Methods). Each protein was represented as an ordered motif string, and TF-IDF-weighted unigram, bigram and trigram profiles were constructed at the AAI50-cluster level. N-grams enriched within individual capsid architecture categories defined recurrent motif cores shared across divergent capsid lineages, indicating that motif-profile organization was associated with broad capsid architecture rather than exclusively lineage-specific variation (Extended Data Figure 4; Methods). At the sequence and embedding-defined group levels, semantic motif profiles provided a higher-resolution view of capsid diversity. For example, HK97-like and picorna-like capsids showed internally stratified subgroups, revealing finer-scale organization within these major capsid architectures (Figure 4b). However, BTV-like capsids represented a notable exception: although they formed distinct latent-space clusters, their semantic motif profiles were more heterogeneous than those of the other architecture categories (Extended Data Figure 5). This discrepancy is consistent with the fact that one-dimensional PLM-derived motif profiles may incompletely capture repeat-rich or helix-dominated architectures in BTV-like capsids, whose defining features depend on repeat register, helix packing, and higher-order structural organization. [26, 27]

**Figure 4.**
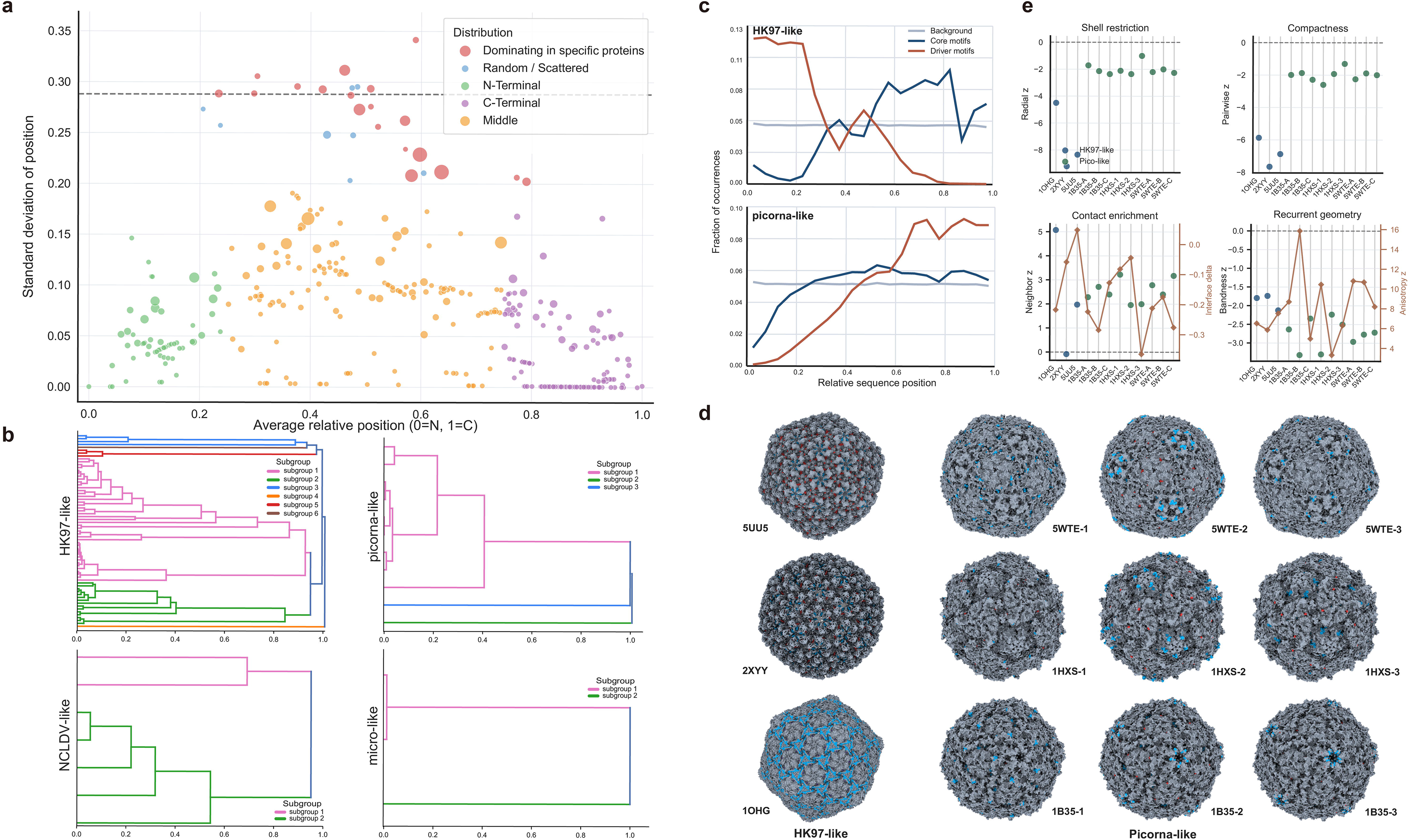
Positional organization of semantic motifs and their structural context across major capsid architectures. (a) Mean relative sequence position and positional standard deviation of semantic motif tokens across capsid proteins. Colours indicate the inferred positional-distribution category, point size represents the mean token fraction, and the dashed horizontal line indicates the expected standard deviation under a uniform positional distribution. (b) Cosine-distance dendrograms of cluster-level semantic motif profiles within the HK97-like, picorna-like, NCLDV-like, and micro-like architecture categories. Each leaf represents an embedding-defined cluster, and branch colours denote subgroups within each architecture category. (c) Relative-position profiles of background token occurrences, conserved core motifs, and lineage-discriminating driver motifs in HK97-like and picorna-like capsid proteins. (d) Surface representations of representative HK97-like assemblies (5UU5, 2XYY, and 1OHG) and picorna-like assemblies (5WTE, 1HXS, and 1B35; chains 1–3), with conserved core-mapped regions highlighted according to their relative sequence positions. The examples are arranged by row as 5UU5 with 5WTE chains 1–3, 2XYY with 1HXS chains 1–3, and 1OHG with 1B35 chains 1–3. (e) Structural properties of the core-mapped regions relative to null residue sets matched for residue count, segment length, and shell radial position. The four panels summarize shell restriction (radial-position z-score), spatial compactness (pairwise-distance z-score), contact enrichment (neighbour-density z-score and interface-fraction difference), and recurrent geometry (bandness and anisotropy z-scores).

Unlike residue-consensus sequence motifs, semantic motifs learned from PLM representations capture recurrent contexts rather than fixed, residue-aligned patterns. At the sequence-position level, most semantic motifs showed characteristic positional preferences along capsid proteins, suggesting that they reflect recurrent positional contexts rather than arbitrary activation fragments (Figure 4a). Within individual capsid architecture categories, semantic motif profiles further resolved lineage-level variation and highlighted conserved structural constraints. For example, HK97-like proteins shared a central semantic-motif signature with greater variability near the N terminus, whereas picorna-like proteins showed increasing motif-profile divergence from the central region toward the N terminus (Figure 4c).

To test whether conserved semantic motif n-gram cores reflect spatial constraints relevant to assembled viral shells, we projected these cores onto representative capsid structures and examined their three-dimensional organization. Conserved motif n-gram cores showed recurrent shell-level localization patterns across three representative HK97-like and picorna-like capsid assemblies. Rather than being diffusely distributed across the protomer or capsomer lattice, core-mapped residues were concentrated in specific structural zones. In HK97-like capsids, core-mapped residues occupied recurrent protomer bands. In picorna-like capsids, they converged on a shared mid-shell band across VP1, VP2, VP3, the three principal picornavirus capsid proteins [28, 29], together with a recurrent shoulder-like feature (Figure 4d). Relative to random residue sets, core-mapped residues were more radially restricted, spatially compact and embedded in denser local structural neighbourhoods. We then used a stricter null model matched on residue count, segment length and shell radial position. Under this matched null, HK97-like core-mapped residues retained the strongest signal for pairwise compactness, whereas picorna-like residues retained both compactness and higher local residue-contact degree (Figure 4e). Core-mapped residues were not consistently enriched at direct inter-subunit interfaces, arguing against their interpretation as generic interface hotspots. Instead, the results are consistent with recurrent scaffold positions and local packing environments associated with capsid-shell geometry, although they do not establish a specific geometry-imposed assembly mechanism. Together, these findings show that SAE-derived semantic motifs are not merely compressed PLM embeddings, but provide an interpretable bridge from capsid sequence representations to conserved shell-level architectural information that is largely invisible to residue-level homology.

### Capsid architectures show broad habitat-associated structuring

To examine the environmental distribution of the recovered capsid repertoire, sample-source metadata were organized into six study-defined major ecosystem categories and their nested sub-ecosystems (see Methods). Among the nested sub-ecosystems, the largest sequence repertoires were recovered from aquatic marine samples (648,272 sequences), followed by aquatic freshwater (523,051), plant-associated habitats (382,937), human-associated samples (259,916), and several extreme-environment systems, including psychrophilic (147,149), thermophilic (87,991) and subsurface/geologic environments (73,987 sequences) (Figure 5a). Sequence-based AAI50 capsid-cluster accumulation curves constructed separately for each major ecosystem did not approach saturation (Extended Data Figure 6), indicating that additional sampling would likely recover further capsid diversity across all ecosystem categories. Sequence-weighted sharing breadth also differed among sub-ecosystems (Figure 5b). Of the 2,701,690 recovered sequences, 2,281,792 (84.5%) were assignable to a major ecosystem category. Among these metadata-annotated sequences, 716,319 (31.4%) belonged to AAI50 capsid clusters observed in only one major ecosystem, whereas 1,565,473 (68.6%) belonged to clusters observed in two or more major ecosystems. Thermophilic environments, wastewater treatment systems and industrial wastewater samples contained particularly high fractions of sequences from cross-ecosystem clusters (78.4%, 78.5% and 86.9%, respectively), whereas plant- and human-associated repertoires had lower shared-sequence fractions (53.0% and 41.4%). Profile-HMM coverage, defined here as the fraction of recovered capsid sequences with at least one significant hit to the reference capsid-profile-HMM set, also differed among sub-ecosystems. Coverage was lower in plant-associated habitats (43.0%) and subsurface environments (46.9%) and higher in aquatic marine (62.2%), aquatic freshwater (60.8%) and animal-associated samples (73.3%) (Figure 5b). These patterns indicate that the expanded capsid repertoire is structured by habitat and is unevenly captured by existing profile HMMs.

**Figure 5.**
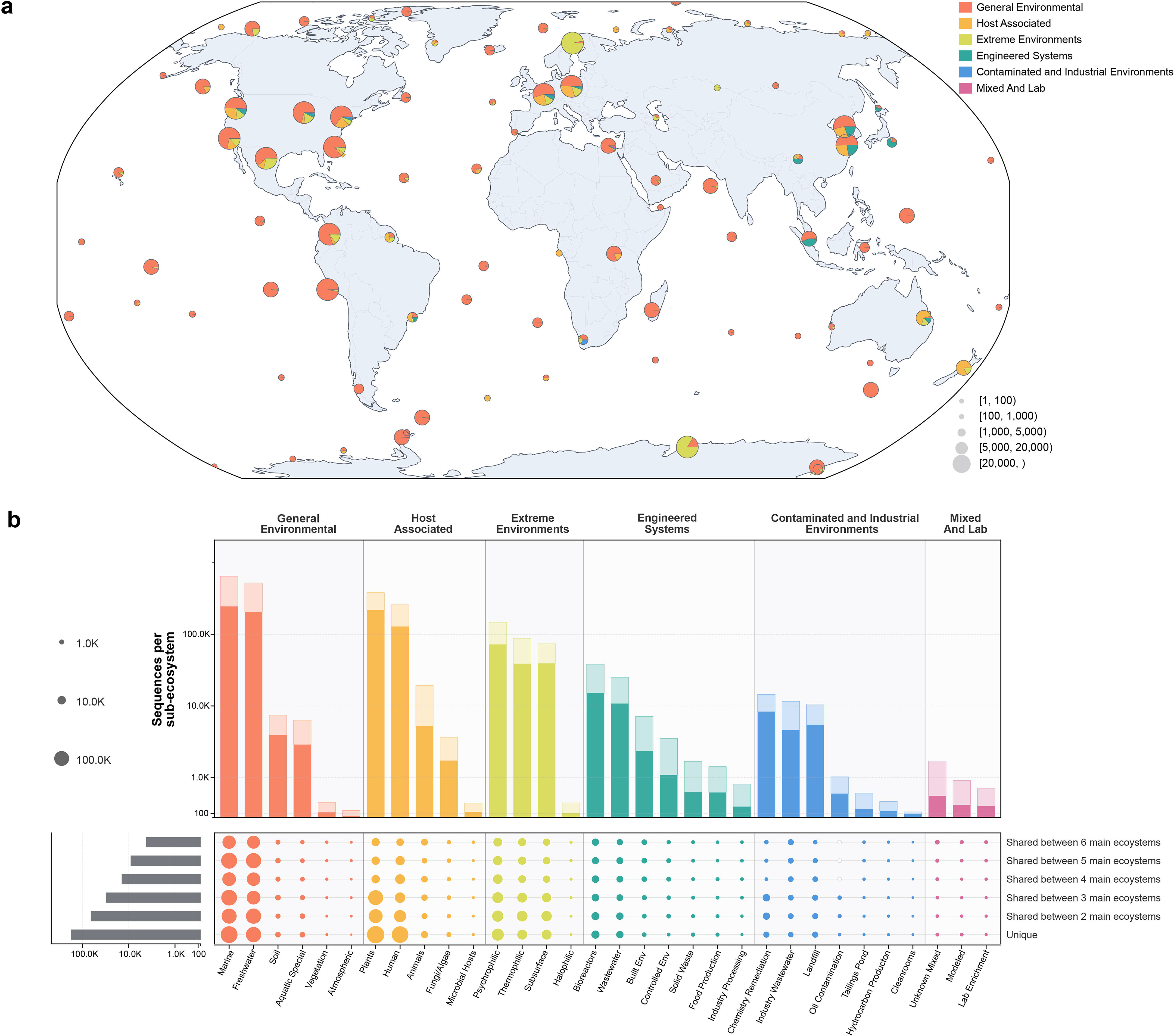
Global biogeographic distribution and ecological sharing breadth of the expanded capsid repertoire. (a) Global distribution of recovered capsid sequences across sampled locations. Sequences were aggregated into 10ř geographic grid cells and displayed as pie markers on a Robinson projection. Pie-marker area is proportional to the total number of recovered sequences in each grid cell, and pie slices indicate the relative contributions of six major ecosystem categories: General Environmental, Host Associated, Extreme Environments, Engineered Systems, Contaminated and Industrial Environments, and Mixed and Lab. (b) Sub-ecosystem-level distribution and cluster-sharing breadth of recovered capsid sequences. Columns represent individual sub-ecosystems grouped by their parent major ecosystem, and the upper bar plot shows the number of recovered sequences assigned to each sub-ecosystem. Rows indicate the ecological breadth of capsid clusters across the six major ecosystem categories, ranging from clusters detected in all six ecosystems to clusters restricted to a single ecosystem. Dot size encodes the number of recovered sequences in each sub-ecosystem contributed by clusters with the corresponding sharing breadth, and the left horizontal bars summarize the total number of clusters in each sharing class. Sequences assigned to unclassified ecosystems were excluded.

Motif-profile-based architecture-category assignments, together with inferred taxonomic and viral genome-type labels, were used to summarize the architectural composition of the expanded repertoire. HK97-like capsids dominated the repertoire (2,324,864 sequences), followed by NCLDV-like (163,695), BTV-like (46,927), micro-like (43,571) and picorna-like capsids (38,637) (Figure 6a). The existing profile-HMM set captured only a fraction of the expanded repertoire, and detection rates varied markedly among capsid architecture categories, ranging from 3.1% in BTV-like and 5.6% in Circoviridae-like capsids to 64.6% in NCLDV-like, 63.2% in micro-like and 61.0% in picorna-like capsids. Sequence redundancy, defined as the mean number of AAI90-dereplicated sequences represented by each AAI50 cluster, also differed among architecture categories. DNA-virus-associated architectures had higher mean AAI50-cluster occupancy than most RNA-virus-associated architectures except levi-like architectures (8.12), with averages of 7.13–14.52 and 2.15–4.55 sequences per cluster, respectively. Together, these comparisons show that different capsid architectures vary substantially in sequence redundancy and profile-HMM coverage, highlighting architecture-specific gaps in current reference models.

**Figure 6.**
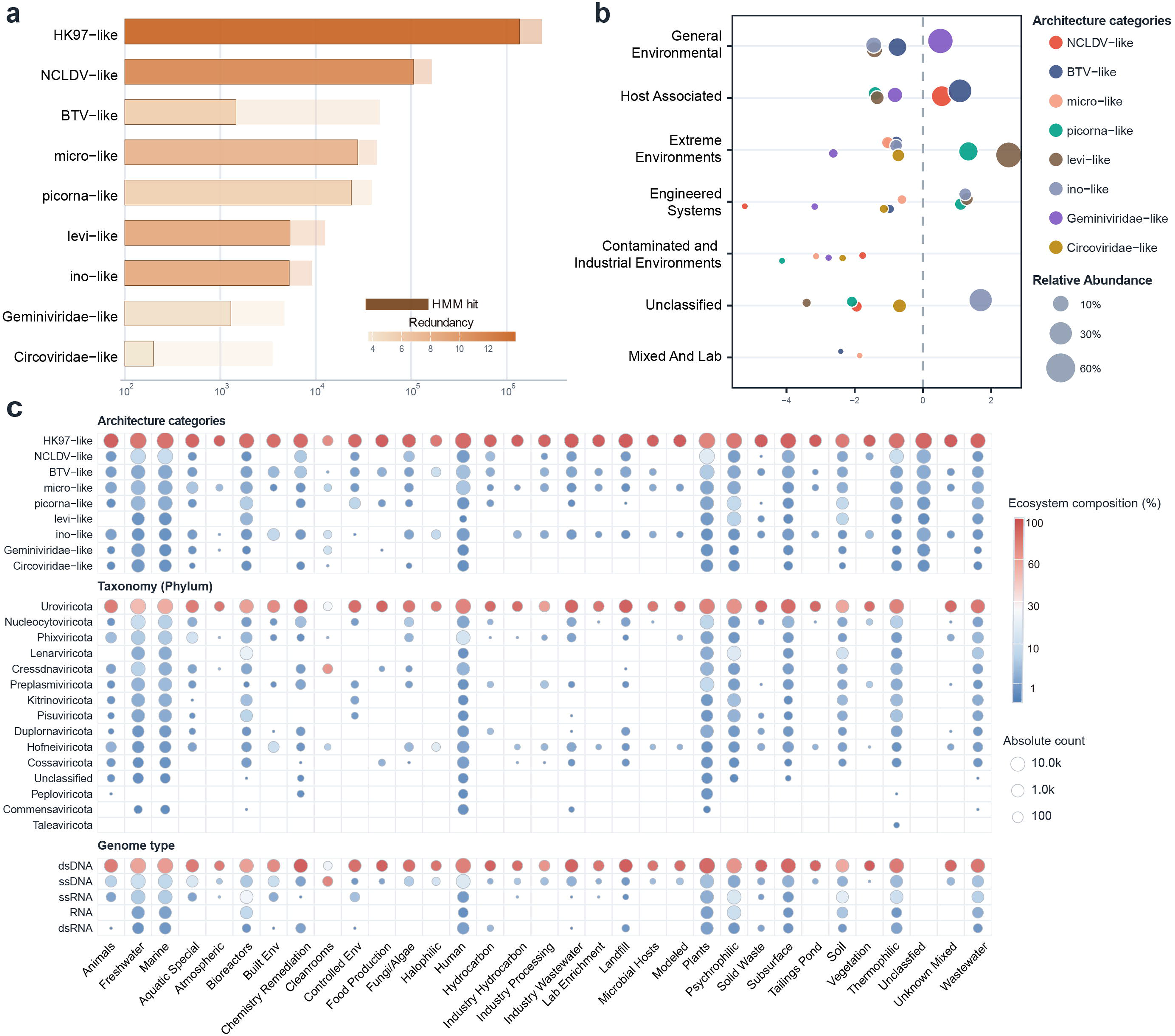
Ecological distribution and enrichment of study-defined capsid architecture categories. (a) Expanded abundance of nine study-defined capsid architecture categories. Horizontal bars show the total expanded member count on a log10 scale, with the darker overlaid segment indicating members supported by Pfam/HMM hits. Bar fill represents redundancy, and labels denote Pfam/HMM-supported counts, with total expanded counts shown in parentheses. (b) Ecosystem-level enrichment of capsid architecture categories across broad environmental categories. Points indicate significant category-environment associations only (q < 0.01 and |log2 enrichment| >= 0.5). The x axis shows log2 enrichment relative to background expectation; positive values indicate enrichment and negative values indicate depletion. Point colour denotes architecture category, and point size represents relative abundance within each category. (c) Sub-ecosystem distribution of capsid signals across three annotation layers: capsid architecture category, taxonomy at the phylum level, and genome type. Each column represents a sub-ecosystem. Point size indicates absolute count, and point colour indicates the percentage contribution of each row category within the corresponding sub-ecosystem. Empty cells indicate no detected counts for that category-sub-ecosystem combination.

Stratifying the repertoire by capsid architecture, inferred genome type and viral lineage revealed additional habitat associations linked to specific capsid architectures (Figure 6b and 6c). For example, RNA-associated capsid signatures, particularly levi-like capsids, showed their highest sequence redundancy in psychrophilic environments. Archaeal-virus-associated lineages were concentrated in high-temperature samples, and Taleaviricota-like signals were detected only in thermophilic environments. The prominence of cressdnavirus-like signals in cleanroom samples and elevated plantvirus-associated diversity in aquatic systems were also observed, which may reflect environmental persistence of small ssDNA virions and hydrological transport of plant-associated viral material, respectively. Together, these analyses reveal habitat-associated differences in the composition of the expanded capsid repertoire at the levels of capsid architecture, inferred genome type and viral lineage.

## Discussion

Environmental viromics has greatly expanded the catalogue of metagenome-derived viral proteins, yet much of this sequence space remains difficult to interpret using primary-sequence homology alone. [6–9] The central challenge is therefore not only to detect divergent viral proteins, but also to place them within biologically meaningful structural and evolutionary contexts. Capsid proteins provide a particularly useful entry point because they are virion-defining components, serve as informative markers of viral diversity, and can retain recognizable foldand architecturelevel relationships after pairwise sequence similarity has become weak or undetectable. [4–6] Here, by integrating capsid-specialized protein language model representations with a semantic-motif framework, we organize divergent metagenome-derived capsids across multiple levels of biological resolution. The resulting framework connects remote capsid sequences to known architecture classes, organizes homology-dark candidates into structurally coherent groups, and links local representation features to recurrent structural environments within representative capsid shells. Rather than simply extending capsid detection, this approach provides an interpretable route from highly divergent sequence space to higher-order capsid organization.

Our benchmark shows that successful binary recognition does not necessarily imply faithful representation of remote or within-capsid diversity. Recent studies using learned protein representations have extended viral protein annotation beyond conventional homology searches and enabled the recovery of highly divergent viral markers. [9, 11, 17] Consistent with this broader potential, all four general-purpose PLMs distinguished capsids from non-capsid proteins accurately under random splits. However, capsid sequences showed higher MLM loss and pseudo-perplexity, and classifier performance declined more sharply under low-identity and family-held-out evaluations. This contrast indicates that broad class separation and remote-family generalization place different demands on protein representations, with the latter requiring features that remain informative when close homologues or entire protein families are withheld from evaluation. ESMCapsid-S and ESMCapsid-C addressed these complementary demands through distinct forms of adaptation. In line with prior evidence that supervised fine-tuning can improve protein-prediction models over frozen pretrained embeddings [9, 20], ESMCapsid-S improved remote capsid detection. By contrast, ESMCapsid-C improved within-capsid representation and increased the linear decodability of taxonomy, capsid architecture category, genome type and host category. Together, these gains show that supervised fine-tuning primarily strengthens remote recognition, whereas continued capsid-domain pretraining provides a representation better suited to organizing biological variation within the capsid family. Beyond remote detection, ESMCapsid-C preserved architecture-associated organization in regions where direct sequence similarity was weak or absent. Previous structure-based studies have shown that viral protein relationships can remain detectable at the fold or architecture level even when sequence similarity is low, and have used structural comparison to connect annotated viral proteins with unannotated families. [5, 6] More broadly, learned protein representations have been shown to encode information related to protein structure across diverse cellular proteins. [12, 14, 15] In our HMM-unsupported embedding-defined groups, predicted structures showed substantially greater similarity within groups than between groups. This separation provides orthogonal evidence that the embedding-defined groups exhibit architecture-associated structural coherence beyond readily detectable sequence homology. ESMCapsid-C therefore extends this general property of protein embeddings to capsid-specialized representation learning, organizing homology-dark capsid candidates into structurally interpretable units that can be compared and prioritized.

Whole-protein embedding groups provide a global map of capsid diversity but do not reveal the local representation features underlying this organization. Building on evidence that sparse autoencoders can separate features stored in superposition within dense PLM embeddings [18, 19], we resolved ESMCapsid-C residue representations into sparse, position-specific semantic motifs that could be localized along sequences and compared across proteins. The resulting motif profiles recapitulated broad architecture-level organization and resolved finer variation within HK97-like and picorna-like capsids, while conserved n-gram cores localized to recurrent structural regions in representative assemblies. Their correspondence with recurrent three-dimensional regions likely reflects structural constraints on sequence evolution and local packing. In capsid proteins, recurrent fold topologies and the geometric requirements of closed-shell assembly repeatedly impose such constraints on local packing [29], providing a basis for the recurrence of SAE-defined cores in comparable structural environments. More broadly, ESM representations have been shown to encode information related to secondary structure, long-range residue contacts, and higher-order sequence dependencies. [12, 14, 16] By decomposing these distributed signals into sparse, position-specific activation patterns, semantic motifs provide a locally interpretable link between the global capsid organization learned by ESMCapsid-C and the structural environments of assembled viral shells.

The environmental analysis indicates that the homology-dark portion of capsid sequence space is habitat-structured rather than uniformly distributed. Previous virome surveys have established habitat as a major axis of viral community organization [2, 7, 30, 31]; our analysis extends this pattern from taxonomic composition to capsid architecture and to the sequence space that remains inaccessible to reference profile-HMM-based annotation. The existing profile-HMM set captured only part of the expanded repertoire, with detection varying across capsid architectures and lower coverage in plant-associated and subsurface repertoires. This indicates that profile-HMM-based surveys may preferentially recover capsid architectures already well represented in reference models, while underrepresenting divergent or previously unmodeled architectures in habitats enriched for such lineages. [9–11] Using ESMCapsid-C-based stratification by capsid architecture, inferred genome type, and viral lineage, we further found habitat associations linked to specific capsid architectures: levi-like capsids showed high sequence redundancy in psychrophilic environments, whereas archaeal-virus-associated lineages were concentrated in high-temperature samples and Taleaviricota-like signals were detected only in thermophilic environments. These patterns suggest that habitat-associated viral partitioning is reflected not only in taxonomic composition, but also in virion architecture. Together, these results show that environmental context shapes both where capsid architectures occur and which parts of capsid diversity remain visible or hidden to conventional profile-HMM annotation.

Our study has several limitations that motivate future work. First, unsupervised clustering of large protein catalogues depends on the choice of clustering hyperparameters, and no single parameterization can optimally capture organization across all levels of viral diversity. As a result, clustering can merge distinct lineages when the threshold is too permissive or split heterogeneous families when it is too stringent. [32] Second, although architecture-category assignments derived from motif profiles are supported here by multiple lines of evaluation, candidate novel architectures and unusual assembly strategies should be regarded as hypotheses generated from computational evidence until experimentally validated. Third, some capsid architectures—particularly BTV-like capsids with repeat-rich or helix-dominated organization—may be less effectively represented by current one-dimensional protein language models, whose representations may not fully capture repeat register, helix packing, and higher-order structural organization. [26, 27] Addressing these limitations will require continued improvement of protein language models, higher-resolution structural analyses, and targeted experimental validation. More broadly, our results suggest that capsid-specialized protein language models provide a framework for tracing structural organization beyond the point at which conventional sequence homology becomes uninformative. As metagenomic sequence space continues to expand, integrating learned representations with structural and experimental virology may therefore make it possible to move from cataloguing viral diversity toward defining the sequence-encoded principles that shape capsid architecture and evolution.

## Methods

### Reference datasets and sequence preprocessing

Protein sequences used for model benchmarking, capsid-specific adaptation, and large-scale environmental screening were obtained from UniProt [33] (release 2025_04), NCBI Virus [34] (accessed September 2025), MGnify Proteins [35] (version 2024_04), and IMG/VR [36] (v4.1; high-confidence genomes only). To reduce redundancy and minimize information leakage, separate MMseqs2 [37] clusterings were generated at 50% amino acid identity for redundancy reduction and at 30% identity to define low-identity-held-out groups. After dereplication and stratified sampling, the binary benchmark contained 5,212 capsid proteins, 40,000 viral non-capsid proteins, and 150,000 cellular proteins, corresponding to capsid-to-viral-non-capsid and capsid-to-cellular-protein ratios of approximately 1:8 and 1:30, respectively. Positive examples for supervised capsid detection comprised proteins explicitly annotated as capsid proteins, coat proteins, major capsid proteins, or shell-forming capsid proteins, supplemented with manually curated annotation terms (Supplementary Note (Additional details of dataset construction)). Sequences containing noncanonical amino acid characters were excluded. Sequences exceeding model-specific context limits were either excluded from analyses requiring full-length proteins or truncated during protein-language-model inference, according to the analysis-specific rules.

For large-scale capsid-protein discovery, the MGnify Proteins and IMG/VR catalogues were combined. Proteins dereplicated at 90% amino acid identity within the source catalogues (AAI90 sequences) were reclustered at 50% identity, yielding 343,633,104 AAI50 clusters from more than 830 million input proteins. One representative sequence from each AAI50 cluster was subjected to model-based screening. For clusters retained as positive, the corresponding AAI90-dereplicated member sequences were carried forward for ecological, structural, and sequence-space-occupancy analyses.

For capsid-specific continued pretraining, we assembled a redundancy-filtered corpus of more than 210,000 capsid proteins from UniProt, NCBI Virus, IMG/VR, and MGnify Proteins. The corpus comprised experimentally supported or curator-annotated capsid proteins together with high-confidence environmental candidates identified during preliminary ESMCapsid-S screening, after the removal of putative non-capsid and cellular contaminants.

### Protein language models and representation extraction

Four pretrained protein language models were benchmarked: ESM2-650M [12], ESM3-open [16], ESMC-600M [38], and ProFluent-E1-600M. [39] Model weights were obtained from the Hugging Face Hub (Supplementary Table 1). For ESMC-based adaptation and downstream analyses, we used the Synthyra/ESMplusplus_large implementation, which provides a Hugging Face-compatible implementation of ESMC. Sequences containing unsupported amino acid characters were removed. Sequences exceeding the model-specific context window were truncated during inference, and no sequence longer than 2,048 model tokens was processed.

For sequence-level analyses, residue-level hidden representations were mean-pooled across amino acid positions after special tokens had been excluded. For token-level analyses, per-residue representations were extracted directly from the specified transformer layer. Unless otherwise stated, model representations were frozen during downstream classifier training. Token-level representations from ESMCapsid-C were subsequently passed to the sparse autoencoder described below.

### Benchmark splits and model evaluation

Model performance was evaluated using three complementary five-fold evaluation schemes: random split, low-identity-held-out split and family-held-out split. In the five-fold random split, the data were divided into five non-overlapping folds; out-of-fold predictions from all five held-out test folds were pooled, and reported metrics were calculated from the pooled confusion matrix. In the low-identity-held-out split, all sequences belonging to the same 30%-amino-acid-identity cluster were assigned to the same fold, thereby reducing leakage from remote homologues. In the family-held-out split, all sequences assigned to the same taxonomic family or curated capsid family were placed in the same fold, such that families represented in a test fold were absent from the corresponding training data. Validation sets were stratified, where compatible with the grouping constraints, by sequence length, source database, and class label. Detailed partitioning and validation procedures are provided in Supplementary Note (Additional details of model training).

Layer-wise representation quality was assessed using frozen-representation probes. For each protein language model, logistic-regression probes were trained on representations extracted from every fourth transformer layer and from each of the final three layers. Performance was summarized using F1 score, precision, recall, and false-positive rate. Model fit to capsid-sequence statistics was evaluated using masked-language-model loss and pseudo-perplexity on held-out capsid and noncapsid proteins.

For comparison with conventional homology-based annotation, the same representative sequences were searched against a capsid-related profile-HMM library using pyhmmer [40, 41] v0.11.3. Unless otherwise stated, hits were retained at an E-value threshold of 1 *×* 10*^−^*^6^ and a minimum domain coverage of 70%. An AAI50 cluster was considered recovered by profile-HMM search when its representative sequence contained at least one hit satisfying both criteria. HMM recovery was evaluated at the AAI50-cluster level to avoid inflation by redundant member sequences.

### Adaptation of ESMCapsid-S and ESMCapsid-C

ESMC was selected as the foundation model for capsid specialization because it provided the best balance between baseline predictive performance and inference efficiency. ESMCapsid-S was derived from ESMC by Low-Rank Adaptation (LoRA) [42] using a supervised contrastive objective [43] for binary capsid detection. Training was performed for 10 epochs with AdamW [44], a learning rate of 1 *×* 10*^−^*^5^, weight decay of 0.01, a batch size of 32, and BF16 mixed precision. Following representation adaptation, lightweight two-layer multilayer perceptron classifiers were trained on mean-pooled hidden representations. Classification thresholds were calibrated on validation data by maximizing F1 subject to a predefined false-positive-rate constraint. Full training and calibration parameters are provided in Supplementary Note (Additional details of model training).

ESMCapsid-C was designed to improve representation quality within capsid sequence space rather than to optimize binary detection alone. Starting independently from the same original ESMC checkpoint used for ESMCapsid-S, all model parameters were updated by continued pretraining on the redundancy-filtered corpus of more than 210,000 capsid sequences. Training used the standard masked-language-model objective [14, 15] with a 15% masking rate for 100,000 optimization steps, using AdamW [44] and a learning rate of 1 *×* 10*^−^*^5^. Held-out capsid sequences were used to monitor masked-language-model loss and pseudo-perplexity, whereas a held-out noncapsid control set was used to assess catastrophic forgetting. Full training details are provided in Supplementary Note (Additional details of model training).

ESMCapsid-C was used for downstream analyses of capsid architecture, taxonomy, ecology, and semantic motifs. To test whether capsid-specific continued pretraining improved biologically informative representations, frozen-representation probes were trained for capsid taxonomy, capsid architecture category, viral genome type, and host category. Key model, screening and sparse-autoencoder settings are summarized in Supplementary Table 2.

### Layer-wise probing and two-stage layer-16 screening

To accelerate metagenomic-scale inference, we implemented a high-stringency, two-stage screening strategy based on layer-16 ESMCapsid-S representations. Layer-wise probes indicated that intermediate ESMCapsid-S representations already separated capsid from non-capsid proteins, supporting the use of layer 16 as a shared representation for large-scale screening. The primary classifier was trained on the standard capsid-versus-non-capsid training set, and its probability threshold was calibrated using an independent validation set. Sequences with a predicted capsid probability below 0.99 were rejected. Sequences meeting the primary threshold were then evaluated by a second classifier trained on a dedicated hard-negative set comprising non-capsid training proteins that were difficult for the primary classifier to distinguish from capsids. Only sequences meeting both the primary-classifier threshold of 0.99 and the hard-negative-classifier threshold of 0.9675 were retained. The hard-negative threshold was calibrated using a separate validation set, and final performance was evaluated on an independent test set that was not used for model training or threshold selection.

### ESMCapsid-C latent-space clustering and group validation

Sequence-level representations generated by ESMCapsid-C were clustered with HDBSCAN [32] to construct a latent-space map of the expanded capsid repertoire. Eight candidate ESMCapsid-C layers (10, 14, 18, 22, 24, 28, 32, and 33) were evaluated across five HDBSCAN parameter combinations. Clustering quality was assessed jointly using mean cluster persistence and the fraction of sequences assigned to noise. Layer 28 provided the best balance between persistence and noise across the parameter scan and was therefore selected for downstream analyses (Extended Data Figure 3). Applying the selected HDBSCAN parameterization to layer-28 representations produced the embedding-defined capsid groups used to analyse repertoire organization, profile-HMM overlap, and taxonomic coherence.

For external taxonomic anchoring, group purity was evaluated using annotated reference sequences assigned to non-noise groups. At each taxonomic rank (realm, kingdom, phylum, and class), sequences lacking an annotation at that rank were excluded. Within each group, the dominant taxon was identified, and group purity was defined as the fraction of annotated sequences assigned to that taxon. Overall purity at each rank was reported as the macro-average, that is, the unweighted mean of group-level dominant-taxon fractions across groups containing valid annotations.

### Sparse-autoencoder training and semantic-motif construction

To obtain interpretable local representations of capsid proteins, sparse autoencoders (SAEs) were trained on residue-level token representations from layer 28 of ESMCapsid-C. Special tokens were excluded, and each residue was represented by a 1,152-dimensional vector. The SAE comparison and final training used the same fixed hard-subset dataset of 150,000 proteins (14,064 BTV-like proteins and 135,936 other capsid proteins; 55,610,678 residue positions). Training tokens were sampled in a sequence-balanced manner by sampling a protein with replacement and then sampling one residue uniformly from that protein. Token vectors were L2-normalized, centred and rescaled using saved global statistics, and clipped at the corresponding per-feature mean *±* 5 standard deviations. Top-k SAEs were evaluated with dictionary sizes *D ∈ {*2,302, 4,604, 11,520, 23,040*}* and sparsity levels *k ∈ {*16, 32, 64*}*. Each model used a linear encoder, rectified linear activations and hard top-k selection [25], followed by linear reconstruction from the selected dictionary vectors. Models were optimized with AdamW [44] using Huber reconstruction loss together with auxiliary residual reconstruction, activation-rate and decoder-norm penalties. The operational *D* = 4,604*, k* = 64 checkpoint was trained for 20 epochs using a configured batch size of 30,240 and was used for all downstream semantic-motif analyses. At the final epoch, its reconstruction loss was 0.1105, explained variance was 0.779 and no dead features were detected. Full preprocessing, initialization, optimization and loss parameters are provided in the Supplementary Note.

A fixed semantic-motif system was constructed from the complete set of 39,001,072 SAE token-activation vectors from 109,220 proteins. After L2 normalization, the 4,604-dimensional activation vectors were zero-padded to 4,608 dimensions and indexed with a FAISS inner-product IndexIVFPQ index [45] comprising 32,768 inverted lists, 64 product-quantization subquantizers and 8 bits per subquantizer. The index was trained on a deterministic blockwise random sample of 2,000,000 token vectors using seed 42. All token vectors were added in batches of 200,000 and searched in batches of 100,000 with nprobe=64 to retrieve 50 neighbours per residue. The resulting weighted graph was partitioned by Leiden community detection [46] at resolution 0.1, producing 364 communities. Each graph vertex, and therefore each residue after special-token removal, received its label directly from its Leiden community. No confidence threshold, rejection rule or unmapped-residue state was used. Raw Leiden identifiers 0–363 were subsequently reordered one-to-one as semantic-motif labels 1–364 according to their positional distributions along proteins; no communities were removed, merged or split.

### Motif strings, motif n-grams, architecture-category assignment, and positional profiling

Each protein was encoded as an ordered residue-wise motif string (*m*_1_*, …, m_L_*), where *m_i_*is the semantic-motif label assigned to residue *i*. All retained residues contributed one label. Consecutive identical labels were retained as separate residue-level tokens, and no run-length compression was applied.

TF-IDF-weighted motif unigram, bigram and trigram profiles were constructed at the AAI50-cluster level from the complete motif strings of member proteins. The resulting profiles were L2-normalized to yield one motif-profile vector per AAI50 cluster. For reference anchoring, 13,504 reference major capsid proteins from NCBI Virus were processed through the same ESMCapsid-C–SAE– semantic-motif pipeline. Reference signatures were derived for eight operational capsid architecture categories: HK97-like, picorna-like, NCLDV-like, BTV-like, Microviridae-like, Geminiviridae-like, Circoviridae-like and levi-like.

Candidate clusters were assigned only when the combined evidence from category-specific core n-gram coverage and global motif-profile similarity exceeded the corresponding category-specific open-set threshold. Cosine similarity and Jensen–Shannon divergence were retained as complementary confidence measures; profile vectors were converted to unit-sum distributions before Jensen– Shannon divergence was calculated.

Positional preferences of motif labels and motif n-grams were summarized along normalized protein coordinates. For an occurrence starting at zero-based position *i* in a protein of length *L*, relative position was defined as *r* = *i/L* and assigned to one of 20 equal-width bins over [0, 1). Background, core and driver profiles were calculated as pooled occurrence frequencies normalized within each profile category, without additional smoothing.

### Structure prediction and validation of homology-dark groups

To test whether embedding-defined groups were structurally coherent beyond detectable sequence homology, five proteins were randomly selected from each eligible HMM-unsupported group included in the structural-validation analysis (22 groups), and their structures were predicted using AlphaFold3. [47] Predicted structures are provided in the Supplementary Materials (20 assigned clusters and 100 structures). All-versus-all pairwise structural alignments were performed with TM-align [48] (4,950 unique structure pairs), and TM-scores were used to quantify structural similarity. Within-group similarity (*n* = 20) was compared with between-group similarity among proteins assigned to the same capsid architecture category (*n* = 39) and with similarity between proteins assigned to different architecture categories (*n* = 151). Statistical significance was assessed using permutation tests (20,000 two-sided label permutations; seed 20260602), with P values adjusted by the Holm procedure. Groups were designated candidate unresolved capsid architectures only when they showed internal structural coherence but lacked confident structural similarity to known capsid structures and known non-capsid homologues according to the predefined criteria.

### Mapping conserved motif n-gram cores onto capsid shells

For the HK97-like and picorna-like lineages, conserved motif n-gram cores were mapped onto experimentally resolved capsid assemblies from the Protein Data Bank [49] or onto AlphaFold3 models of the corresponding assemblies when experimental structures were unavailable. Three representative assemblies from each architecture category were analysed (Supplementary Table 3). The particle centre was defined as the mean coordinate of all C*α* atoms in the assembly. For each residue, normalized radial position was calculated as (||x - c|| - r_min)/(r_max - r_min), where x is the residue C*α* coordinate, c is the particle centre, and r_min and r_max are the minimum and maximum assembly-wide C*α* radii, respectively. Residues were classified as belonging to the inner, middle, or outer shell when their normalized radial positions were <0.33, 0.33 to <0.67, or *≥*0.67, respectively. Conserved motif cores were mapped back to the corresponding sequence positions, intersected with structure-resolved residues in the representative chain, and merged into contiguous ranges. Motif-to-structure projection, matched null models, compactness measures, and inter-subunit interface analyses are described in Supplementary Note (Additional details of sparse feature analyses).

### Ecological metadata harmonization and diversity analyses

Sample-source metadata associated with MGnify Proteins and IMG/VR proteins were retrieved from the corresponding sample-level records. Where available, source-environment descriptors were mapped to the five-level GOLD ecosystem-classification hierarchy—Ecosystem, Ecosystem Category, Ecosystem Type, Ecosystem Subtype, and Specific Ecosystem—which was developed to standardize the classification of metagenomic samples. [50, 51] To harmonize heterogeneous metadata vocabularies across MGnify and IMG/VR and enable consistent cross-database comparisons, the original GOLD classification paths and database-specific sample-source terms were reorganized into six broader major ecosystem categories and their constituent sub-ecosystems. The resulting categories were used for the ecological summaries below. Sequences lacking sufficient source metadata were retained as Unclassified. Sequence-level taxonomy, viral genome type, and related biological labels were assigned using ESMCapsid-C representation-based prediction models.

An AAI50 capsid cluster was classified as single-ecosystem when all metadata-annotated member sequences originated from the same major ecosystem category and as cross-ecosystem when annotated members occurred in two or more major ecosystem categories. Members lacking ecosystem metadata did not contribute to this classification. Sequence-level accumulation curves were generated separately for each major ecosystem by randomly subsampling metadata-annotated sequences without replacement and counting the number of unique AAI50 capsid clusters recovered. For each ecosystem, 20 logarithmically spaced subsample sizes, ranging from 50 sequences to the total number of available sequences, were evaluated. Each subsample size was repeated 500 times. Mean cluster recovery and central 95% subsampling intervals were summarized using the 2.5th and 97.5th percentiles of the repeated subsamples. Sequence redundancy within each architecture category was defined as the mean number of AAI90-dereplicated sequences represented by an AAI50 cluster.

### Statistical analysis and reproducibility

Unless otherwise stated, statistical analyses were performed in Python 3.11. Model training and inference used PyTorch 2.1.1. [52] Sequence clustering used MMseqs2 release 18 [37], and profile-HMM searches used pyhmmer v0.11.3. [40, 41] Figures were generated using matplotlib, seaborn, and plotly. All tests were two-sided unless otherwise specified. Independent distributional comparisons used Wilcoxon rank-sum tests. Structural-coherence analyses used permutation tests with Holm adjustment. For analyses involving multiple feature- or motif-level comparisons, P values were adjusted using the Benjamini–Hochberg procedure [53], unless the Holm procedure [54] was explicitly specified. Confidence intervals were estimated by bootstrap resampling as specified for each analysis. [55] Model training and inference were conducted on a high-performance computing cluster equipped with 128 CPU cores, 768 GB RAM, and NVIDIA A800 or L40 GPUs.

## Data availability

Protein sequences and annotations analysed in this study were obtained from UniProt (release 2025_04), NCBI Virus (accessed September 2025), MGnify Proteins (version 2024_04) and IMG/VR (v4.1; high-confidence genomes only), as described in the Methods. Taxonomic labels were standardized using the ICTV 2024 Master Species List, MSL40 release v2, and host labels were obtained from Virus-Host DB release 231.

The datasets generated and analysed during this study are available in Figshare at https://doi.org/10.6084/m9.figshare.33453790. Source Data files underlying all panels of Figures 2–6 and Extended Data Figures 1–6 are available in Figshare at https://doi.org/10.6084/m9.figshare.33453826. The ESMCapsid-S and ESMCapsid-C model weights are available from the Hugging Face Hub at https://huggingface.co/Shuofang127/ESMCapsid-S and https://huggingface.co/Shuofang127/ESMCapsid-C, respectively.

## Code availability

All code used to train and evaluate ESMCapsid, perform large-scale capsid screening and downstream analyses, and generate the figures and tables is available at https://github.com/liusihang/ESMCapsid. This repository includes the model-training, benchmarkevaluation, sparse-autoencoder and semantic-motif analysis, structural-analysis, inference-pipeline, and figure-generation code used in this study.

## Acknowledgements

This study was funded by the National Natural Science Foundation of China (Grant No. 52570110). The views presented here do not represent those of the sponsors. We also acknowledge the Scientific Computing Service Platform of Tongji University for providing computational resources.

## Author contributions

Sihang Liu conceived the study, developed the methodology and models, performed all computational analyses, curated the data, prepared the figures and visualizations, and wrote the original draft. Pingfeng Yu reviewed and edited the manuscript. Hong Wang supervised the study, acquired funding, and reviewed and edited the manuscript. Siqing Xia supervised the study and acquired funding. All authors reviewed and approved the final manuscript.

## Extended Data legends

**Extended Data Figure 1.**
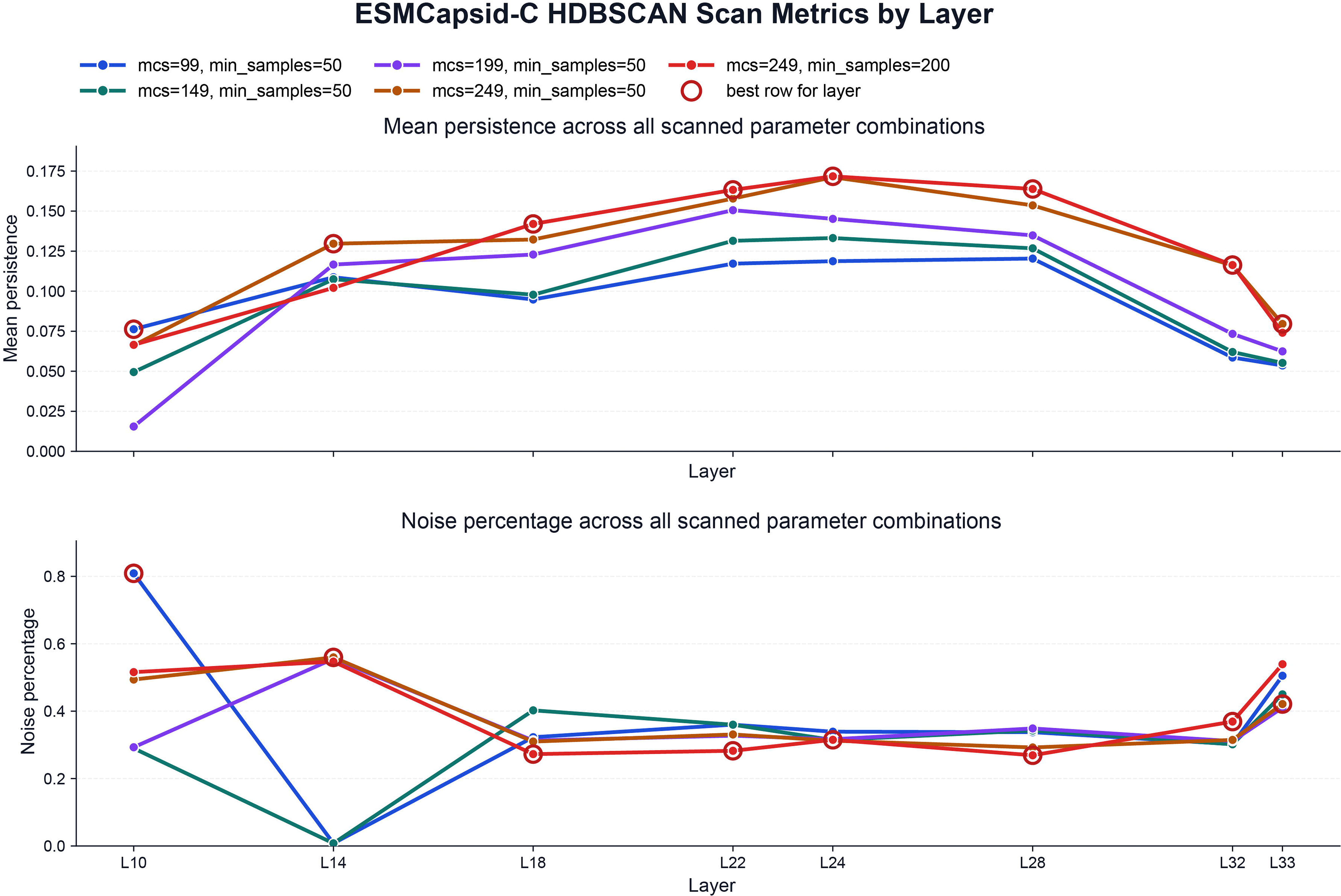
Binary t-SNE visualization of capsid and other proteins across four generic protein language models. The four panels show t-SNE projections of sequence embeddings from ESMC-600M, ESM3-open, ESM2-650M and ProFluent-E1-600M for the binary benchmark. Capsid proteins are shown separately from other proteins, where the latter group includes cellular proteins and viral proteins not annotated as capsids.

**Extended Data Figure 2.**
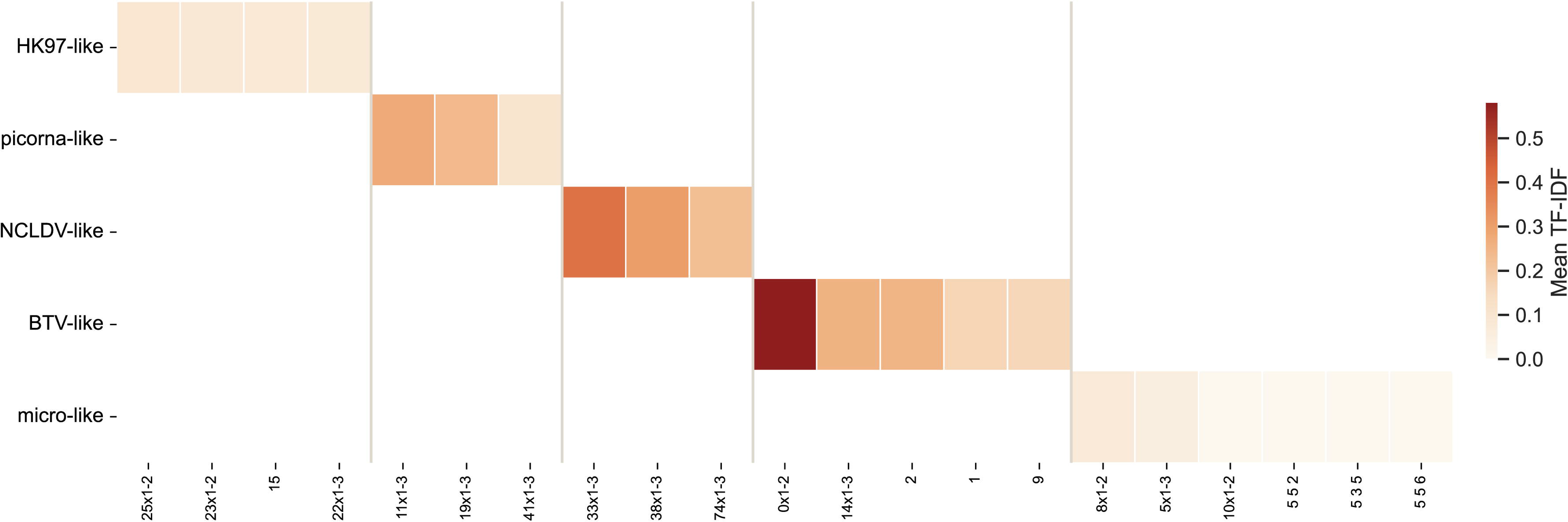
Layer-wise linear-probe performance of ESMCapsid-S and final Layer16 two-head screening performance. Sequence-level mean-pooled ESMC representations from each layer were evaluated using a single-head linear probe on the capsid versus non-capsid benchmark. For each layer, bars show the resulting F1 score, accuracy, and recall. The gray line indicates the expected inference-time trend across layers. The final column shows the performance of the final Layer16 two-step screening scheme, in which a shared Layer16 representation is first scored by the normal head and then filtered by the hard-negative head.

**Extended Data Figure 3.**
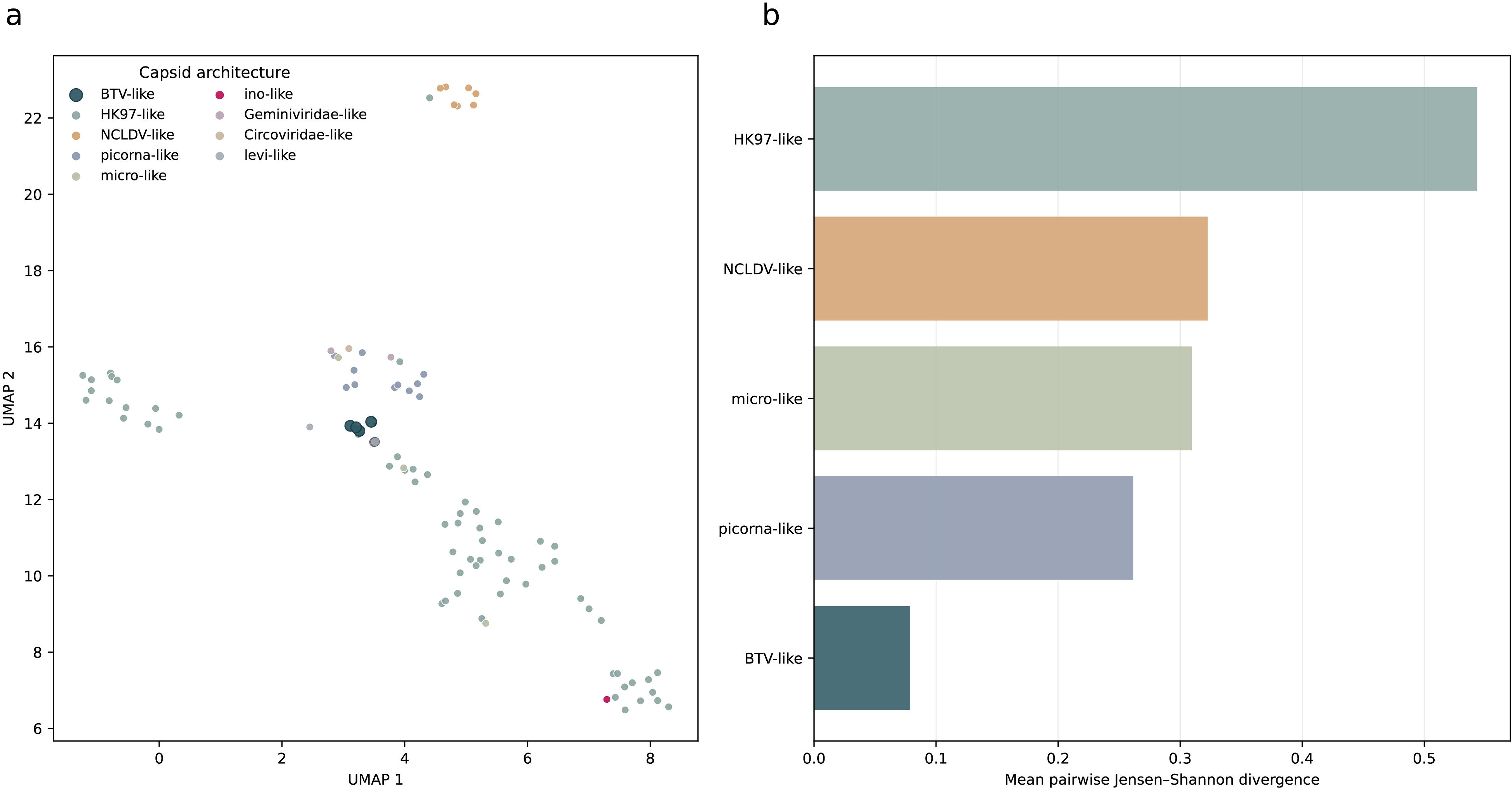
Layerand parameter-scan comparison for HDBSCAN clustering of sequence-level ESMCapsid-C embeddings. Eight candidate embedding layers (10, 14, 18, 22, 24, 28, 32, and 33) were evaluated across five HDBSCAN parameter combinations (min_cluster_size/min_samples: 99/50, 149/50, 199/50, 249/50, and 249/200). The upper panel shows mean cluster persistence and the lower panel shows the fraction of sequences assigned to noise for each layer-parameter combination. Red circles mark the best-performing parameter setting selected within each layer.

**Extended Data Figure 4.**
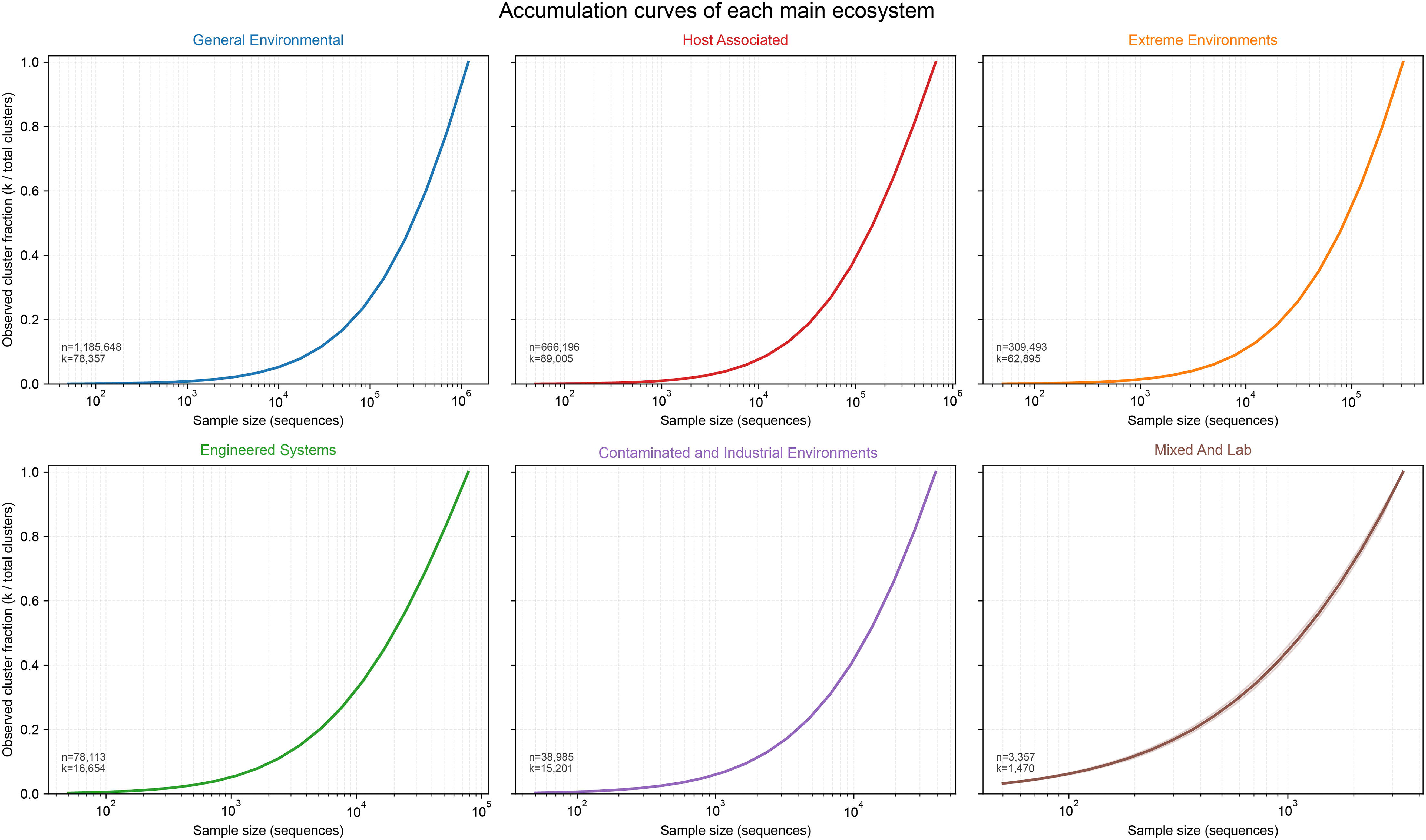
Semantic-motif grammar across major capsid architecture categories. Rows denote study-defined architecture categories and columns denote merged motif families. Repeated unigram, bigram and trigram forms of the same motif token were collapsed for display. Cell colour represents the mean TF-IDF weight within an architecture category; blank cells indicate that the motif family was not retained among the eight highest-weight recurrent core features for that category. Only categories with multi-cluster support are shown.

**Extended Data Figure 5.**
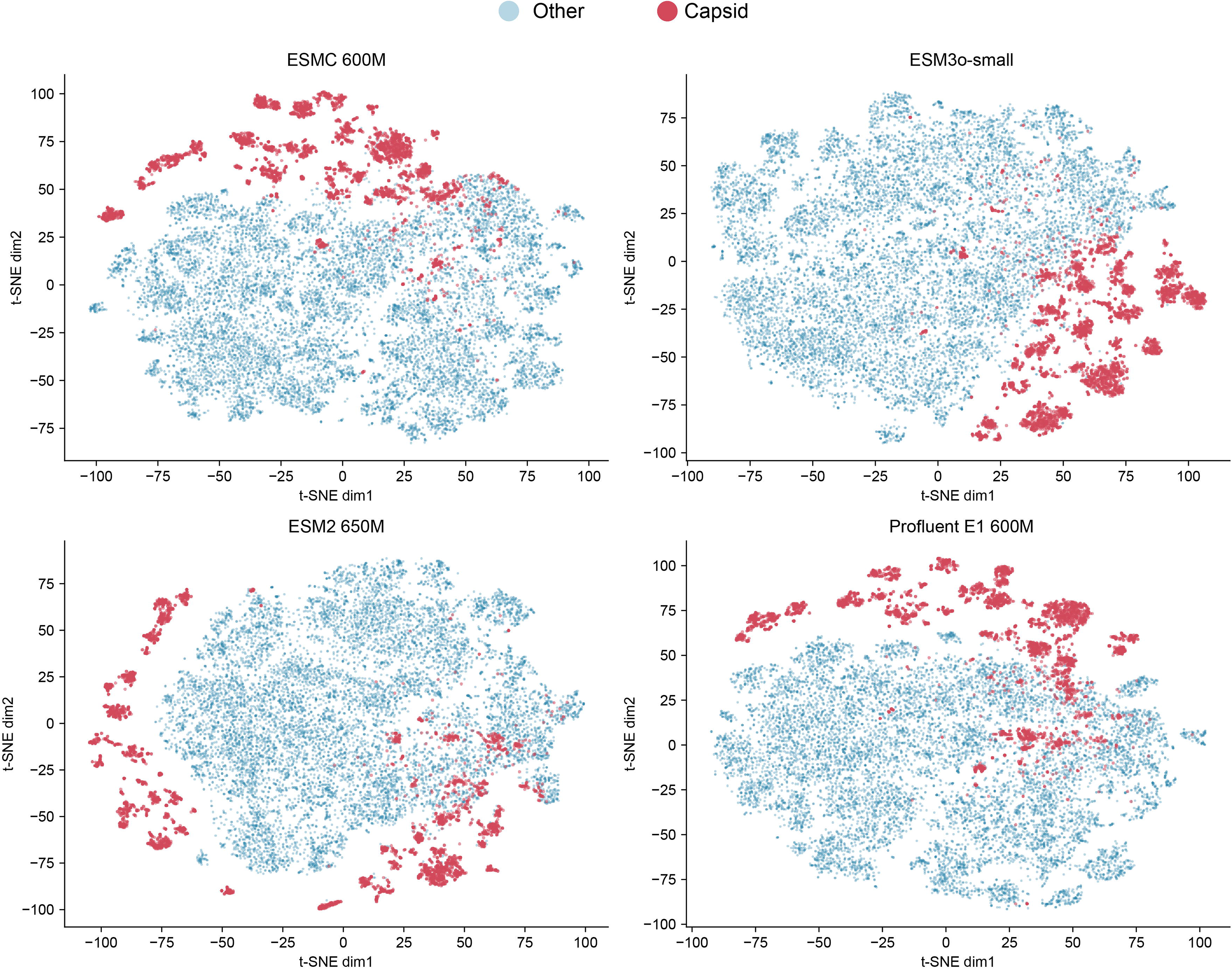
BTV-like capsids show localized latent-space placement but elevated motif-profile heterogeneity. a, UMAP projection of cluster-level motif-profile embeddings under the current architecture grouping. BTV-like clusters are emphasized. b, Positional-profile divergence across architecture categories. For motifs present in at least 80% of clusters in a category, divergence was calculated as the mean pairwise Jensen-Shannon divergence between 20-bin positional histograms. The right-hand values show the fraction of evaluable motifs with divergence greater than 0.16. BTV-like is represented by two retained clusters in this analysis.

**Extended Data Figure 6.**
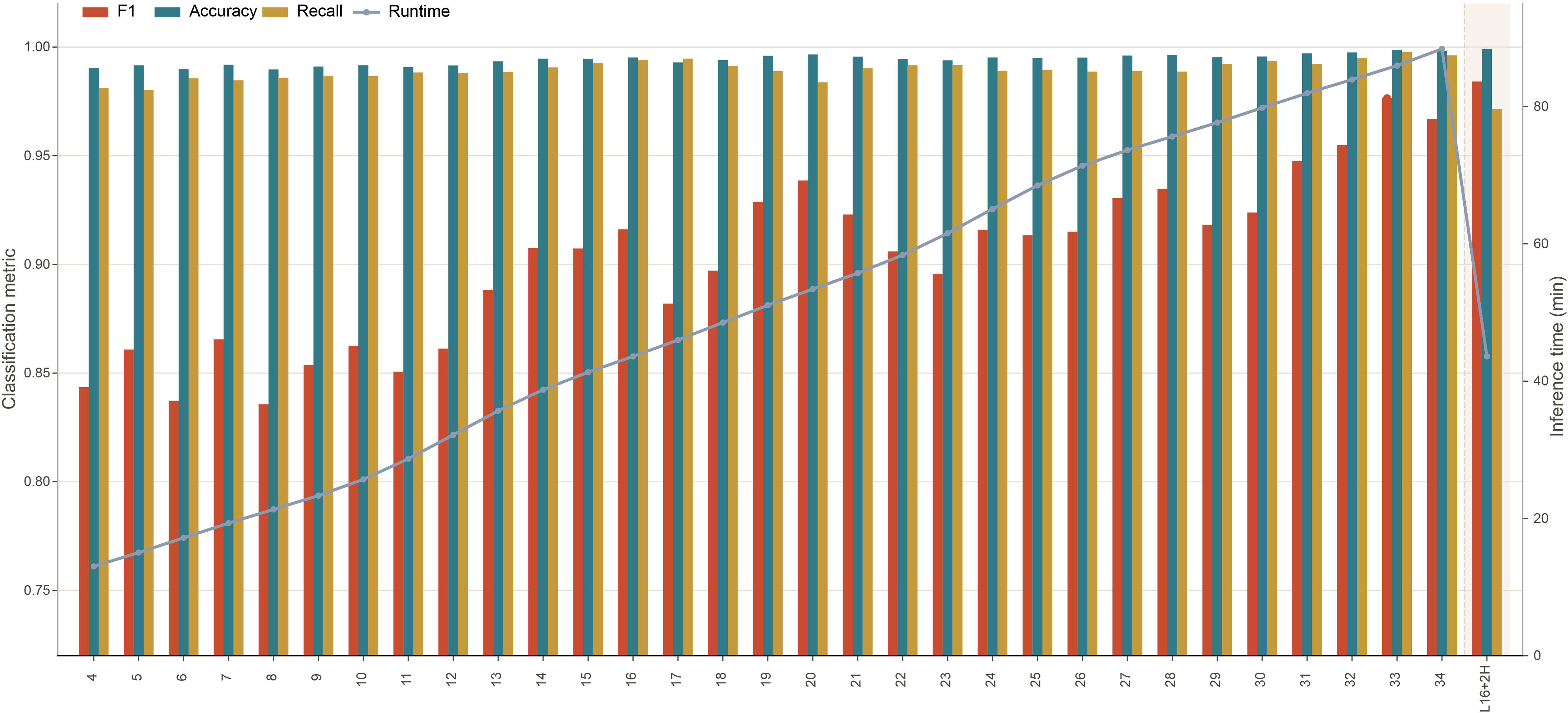
Sequence-level capsid-cluster accumulation curves across major ecosystem categories. Within each assigned major ecosystem, metadata-annotated sequences were repeatedly subsampled without replacement at 20 logarithmically spaced sequence counts, from 50 sequences to the full ecosystem-specific sequence set. Each sequence count was evaluated in 500 replicates. Curves show the mean fraction of ecosystem-specific AAI50 capsid clusters recovered, and shaded bands show the central 95% subsampling interval. Insets report the total number of sequences (n) and AAI50 capsid clusters (k). The sampling unit is sequence count; these curves do not estimate accumulation under newly collected independent environmental samples.

## References

[1] Suttle, C. A. Viruses in the sea. Nature 437, 356–361 (2005).

[2] Brum, J. R. et al. Patterns and ecological drivers of ocean viral communities. Science 348, 1261498 (2015).

[3] International Committee on Taxonomy of Viruses Executive Committee, et al. The new scope of virus taxonomy: partitioning the virosphere into 15 hierarchical ranks. Nat. Microbiol. 5, 668–674 (2020).

[4] Krupovic, M., Dolja, V. V. & Koonin, E. V. Origin of viruses: primordial replicators recruiting capsids from hosts. Nat. Rev. Microbiol. 17, 449–458 (2019).

[5] Abrescia, N. G. A. et al. Structure unifies the viral universe. Annu. Rev. Biochem. 81, 795–822 (2012).

[6] Nomburg, J. et al. Birth of protein folds and functions in the virome. Nature 633, 710–717 (2024).

[7] Paez-Espino, D., et al. Uncovering Earth’s virome. Nature 536, 425–430 (2016).

[8] Pavlopoulos, G. A. et al. Unraveling the functional dark matter through global metagenomics. Nature 622, 594–602 (2023).

[9] Flamholz, Z. N., Biller, S. J. & Kelly, L. Large language models improve annotation of prokaryotic viral proteins. Nat. Microbiol. 9, 537–549 (2024).

[10] Hamamsy, T. et al. Protein remote homology detection and structural alignment using deep learning. Nat. Biotechnol. 42, 975–985 (2024).

[11] Liu, W. et al. PLMSearch: protein language model powers accurate and fast sequence search for remote homology. Nat. Commun. 15, 2775 (2024).

[12] Lin, Z. et al. Evolutionary-scale prediction of atomic-level protein structure with a language model. Science 379, 1123–1130 (2023).

[13] van Kempen, M. et al. Fast and accurate protein structure search with Foldseek. Nat. Biotechnol. 42, 243–246 (2024).

[14[ Rives, A., et al. Biological structure and function emerge from scaling unsupervised learning to 250 million protein sequences. Proc. Natl Acad. Sci. USA 118, e2016239118 (2021).

[15] Elnaggar, A. et al. ProtTrans: towards understanding the language of life through selfsupervised learning. IEEE Trans. Pattern Anal. Mach. Intell. 44, 7112–7127 (2022).

[16] Hayes, T. et al. Simulating 500 million years of evolution with a language model. Science 387, 850–858 (2025).

[17] Hou, X. et al. Using artificial intelligence to document the hidden RNA virosphere. Cell 187, 6929–6942.e16 (2024).

[18] Gujral, O., Bafna, M., Alm, E. & Berger, B. Sparse autoencoders uncover biologically interpretable features in protein language model representations. Proc. Natl Acad. Sci. USA 122, e2506316122 (2025).

[19] Simon, E. & Zou, J. InterPLM: discovering interpretable features in protein language models via sparse autoencoders. Nat. Methods 22, 2107–2117 (2025).

[20] Schmirler, R., Heinzinger, M. & Rost, B. Fine-tuning protein language models boosts predictions across diverse tasks. Nat. Commun. 15, 7407 (2024).

[21] van der Maaten, L. & Hinton, G. Visualizing data using t-SNE. J. Mach. Learn. Res. 9, 2579– 2605 (2008).

[22] Altschul, S. F. et al. Basic local alignment search tool. J. Mol. Biol. 215, 403–410 (1990).

[23] Hua, J. et al. Capsids and genomes of jumbo-sized bacteriophages reveal the evolutionary reach of the HK97 fold. mBio 8, e01579–17 (2017).

[24] Yang, Y. et al. Capsid structure of bacteriophage ΦKZ provides insights into assembly and stabilization of jumbo phages. Nat. Commun. 15, 6551 (2024).

[25] Gao, L. et al. Scaling and evaluating sparse autoencoders. International Conference on Learning Representations (2025).

[26] Grimes, J. M. et al. The atomic structure of the bluetongue virus core. Nature 395, 470–478 (1998).

[27] Zhang, Q. et al. The structure of a 12-segmented dsRNA reovirus: new insights into capsid stabilization and organization. PLoS Pathog. 19, e1011341 (2023).

[28] Rossmann, M. G. et al. Structure of a human common cold virus and functional relationship to other picornaviruses. Nature 317, 145–153 (1985).

[29] Roos, W. H., Bruinsma, R. & Wuite, G. J. L. Physical virology. Nat. Phys. 6, 733–743 (2010).

[30] Gregory, A. C. et al. Marine DNA viral macro- and microdiversity from pole to pole. Cell 177, 1109–1123.e14 (2019).

[31] Ma, B. et al. Biogeographic patterns and drivers of soil viromes. Nat. Ecol. Evol. 8, 717–728 (2024).

[32] McInnes, L., Healy, J. & Astels, S. hdbscan: hierarchical density based clustering. J. Open Source Softw. 2, 205 (2017).

[33] The UniProt Consortium. UniProt: the Universal Protein Knowledgebase in 2025. Nucleic Acids Res. 53, D609–D617 (2025).

[34] Brister, J. R., Ako-adjei, D., Bao, Y. & Blinkova, O. NCBI Viral Genomes Resource. Nucleic Acids Res. 43, D571–D577 (2015).

[35] Richardson, L. et al. MGnify: the microbiome sequence data analysis resource in 2023. Nucleic Acids Res. 51, D753–D759 (2023).

[36] Camargo, A. P. et al. IMG/VR v4: an expanded database of uncultivated virus genomes within a framework of extensive functional, taxonomic, and ecological metadata. Nucleic Acids Res. 51, D733–D743 (2023).

[37] Steinegger, M. & Söding, J. MMseqs2 enables sensitive protein sequence searching for the analysis of massive data sets. Nat. Biotechnol. 35, 1026–1028 (2017).

[38] Candido, S. et al. Language modeling materializes a world model of protein biology. Preprint at bioRxiv (2026); doi:10.64898/2026.06.03.729735.

[39] Jain, S., et al. E1: retrieval-augmented protein encoder models. Preprint at bioRxiv (2025); doi:10.1101/2025.11.12.688125.

[40] Eddy, S. R. Accelerated profile HMM searches. PLoS Comput. Biol. 7, e1002195 (2011).

[41] Larralde, M. & Zeller, G. PyHMMER: a Python library binding to HMMER for efficient sequence analysis. Bioinformatics 39, btad214 (2023).

[42] Hu, E. J. et al. LoRA: low-rank adaptation of large language models. International Conference on Learning Representations (2022).

[43] Khosla, P., et al. Supervised contrastive learning. Advances in Neural Information Processing Systems 33, 18661–18673 (2020).

[44] Loshchilov, I. & Hutter, F. Decoupled weight decay regularization. International Conference on Learning Representations (2019).

[45] Johnson, J., Douze, M. & Jégou, H. Billion-scale similarity search with GPUs. *IEEE Trans*. Big Data 7, 535–547 (2021).

[46] Traag, V. A., Waltman, L. & van Eck, N. J. From Louvain to Leiden: guaranteeing well-connected communities. Sci. Rep. 9, 5233 (2019).

[47] Abramson, J. et al. Accurate structure prediction of biomolecular interactions with AlphaFold 3. Nature 630, 493–500 (2024).

[48] Zhang, Y. & Skolnick, J. TM-align: a protein structure alignment algorithm based on the TM-score. Nucleic Acids Res. 33, 2302–2309 (2005).

[49] wwPDB Consortium. Protein Data Bank: the single global archive for 3D macromolecular structure data. Nucleic Acids Res. 47, D520–D528 (2019).

[50] Ivanova, N. et al. A call for standardized classification of metagenome projects. Environ. Microbiol. 12, 1803–1805 (2010).

[51] Mukherjee, S. et al. Twenty-five years of Genomes OnLine Database (GOLD): data updates and new features in v.9. Nucleic Acids Res. 51, D957–D963 (2023).

[52] Paszke, A. et al. PyTorch: an imperative style, high-performance deep learning library. Advances in Neural Information Processing Systems 32, 8024–8035 (2019).

[53] Benjamini, Y. & Hochberg, Y. Controlling the false discovery rate: a practical and powerful approach to multiple testing. J. R. Stat. Soc. B 57, 289–300 (1995).

[54] Holm, S. A simple sequentially rejective multiple test procedure. Scand. J. Stat. 6, 65–70 (1979).

[55] Efron, B. Bootstrap methods: another look at the jackknife. Ann. Stat. 7, 1–26 (1979).

