## Supplementary Information for "Capsid-specialized protein language models reveal higher-order viral architecture from sequence"

### Supplementary Note

#### Additional details of dataset construction

##### Dataset curation, redundancy control and benchmark splits

Protein sequences used for model benchmarking and capsid-specific adaptation were obtained from the UniProt, NCBI Virus, MGnify Proteins and IMG/VR resources described in the main Methods. Candidate positive examples were retained when their annotations explicitly identified a capsid protein, coat protein, major capsid protein or shell-forming capsid protein, together with a manually curated set of synonymous capsid terms. Proteins annotated as portal, terminase, tail, tail fibre, baseplate, tape-measure protein, envelope, matrix, spike, glycoprotein, packaging ATPase, scaffold or prohead protease were excluded from the positive set unless independent evidence supported a capsid assignment. Viral proteins not assigned as capsids were used as challenging viral negatives, and cellular proteins from UniProt were used as cellular negatives.

Sequences containing unsupported amino-acid characters were removed. During protein-language-model inference, sequences were limited to 2,048 model tokens after special-token accounting. The labelled benchmark contained 5,212 capsid proteins, 40,000 viral non-capsid proteins and 150,000 cellular proteins. MMseqs2 was used at 50% amino-acid identity for redundancy control and at 30% identity to construct low-identity evaluation groups. All members of an identity cluster were assigned to the same fold. Three five-fold evaluation schemes were used: a stratified random split, a 30%-identity cluster-held-out split and a family-held-out split. For family-held-out evaluation, only valid family-level labels ending in -viridae with at least 20 positive proteins were retained, and each eligible family was withheld in turn while the negative partition was kept fixed.

#### Additional details of model training

##### Protein language model benchmark and representation-probe settings

Four generic protein language models were benchmarked: ESM2-650M, ESM3-open, ESMC-600M and ProFluent-E1-600M (Supplementary Table 1). Frozen residue-level representations were extracted from every fourth transformer layer and from each of the final three layers. Special tokens were excluded before mean pooling to obtain sequence-level representations. Logistic-regression probes were trained for capsid-versus-non-capsid classification under the random, low-identity-held-out and family-held-out schemes. Performance was summarized using precision, recall, F1 score, balanced accuracy and false-positive rate. Two-dimensional t-SNE projections were used only for visualization and did not contribute to classifier training or model selection.

The benchmark comparisons were performed on the same fixed sequence partitions for all models. For the random split, out-of-fold predictions from the five held-out folds were pooled and metrics were calculated from the pooled confusion matrix. For low-identity-held-out and family-held-out schemes, test metrics were calculated independently in each held-out fold or taxon and summarized across the corresponding held-out units. The model checkpoints, model sizes and representation outputs used in this study are summarized in Supplementary Table 1.

#### Masked-language-model loss and pseudo-perplexity

Masked-language-model loss was calculated by masking 15% of amino-acid positions in each evaluation pass and averaging the negative log probability assigned to the true residue over masked positions only:

$$L_{\mathrm{MLM}} = -\frac{1}{|M|}\sum_{i\in M}\log p(x_i \mid x_{\setminus M})$$

Here, $x$ denotes the protein sequence, $M$ the set of masked residue positions and $x_{\setminus M}$ the masked input sequence. Pseudo-perplexity was calculated by masking one residue at a time and summing the conditional log likelihood across the full sequence:

$$\mathrm{PPPL}(x) = \exp\left[-\frac{1}{L}\sum_{i=1}^{L}\log p(x_i \mid x_{\setminus i})\right]$$

Here, L is the sequence length after special-token removal. Lower masked-language-model loss and lower pseudo-perplexity indicate better fit to the corresponding sequence distribution. Capsid and non-capsid distributions were compared using two-sided Wilcoxon rank-sum tests.

#### ESMCapsid-S adaptation, frozen probes and layer-16 two-stage screening

ESMCapsid-S was initialized from ESMC and adapted for binary capsid detection by Low-Rank Adaptation with a supervised contrastive objective. Training used AdamW for 10 epochs with a learning rate of $1 \times 10^{-5}$, weight decay of 0.01, a batch size of 32 and BF16 mixed precision. After representation adaptation, frozen sequence representations were classified with two-layer multi-layer perceptron heads with hidden widths of 128 and 256, ReLU activation and dropout of 0.1. The same probe architecture was used for the frozen-representation annotation tasks described in Supplementary Note (Additional details of model training), with task-specific labels and class weighting.

For metagenomic-scale inference, layer-wise probe performance and recorded inference time were evaluated across ESMCapsid-S layers (Extended Data Figure 2). Layer 16 was selected as the shared representation for a two-stage screen. A normal head first retained sequences with capsid probability at least 0.99. Retained sequences were then evaluated by a hard-negative head, and sequences with hard-negative-head probability at least 0.9675 were kept. Thresholds were fixed from independent validation sets before final test evaluation. Key adaptation and inference settings are summarized in Supplementary Table 2.

##### ESMCapsid-C continued pretraining and biological annotation probes

ESMCapsid-C was initialized independently from the same original ESMC checkpoint and continued-pretrained on a redundancy-filtered corpus of more than 210,000 capsid proteins from UniProt, NCBI Virus, IMG/VR and MGnify Proteins. All model parameters were updated using the standard masked-language-model objective with a 15% masking rate for 100,000 optimization steps. Training used AdamW with a learning rate of $1 \times 10^{-5}$. Held-out capsid proteins were used to monitor domain adaptation, and a held-out non-capsid set was used to monitor catastrophic forgetting.

For downstream annotation probes, reference capsid proteins were dereplicated at 50% amino-acid identity. Taxonomic labels were standardized against the ICTV 2024 Master Species List, MSL40 release v2. Host labels were obtained by matching viral taxonomic identifiers to Virus-Host DB release 231. Architecture-category and genome-type labels were derived from standardized taxonomy and cross-checked against structural literature where available. Layer-wise evaluation identified layer 33 for taxonomy, architecture category, genome type and host-category prediction, whereas layer 28 was used for the clustering and token-level sparse-autoencoder analyses. Class-weighted losses were used for imbalanced annotation tasks.

#### Additional details of sparse feature analyses

##### Sparse-autoencoder training, token-graph community detection and motif-string construction

**SAE training data and residue sampling** Residue-level representations were extracted from layer 28 of ESMCapsid-C. Special tokens were removed, leaving one 1,152-dimensional representation for each amino acid residue. The SAE model comparison and final training used the same fixed hard-subset dataset used for the April 2026 retraining analysis. This dataset comprised 150,000 proteins, including 14,064 BTV-like proteins and 135,936 other capsid proteins, and contained 55,610,678 residue positions before epoch-level sampling.

Training examples were generated using sequence-balanced rather than residue-length-weighted sampling. For each of 4,000,000 draws in an epoch, a protein was sampled uniformly with replacement from the fixed protein set, after which one residue was sampled uniformly from the selected protein. Thus, long proteins did not contribute proportionally more training examples solely because of their length. Sampling used a base random seed of 42, and the seed for epoch e was 42 + 1,000e. Token vectors were L2-normalized. Global per-feature centring and scaling statistics were estimated from the preprocessed epoch chunks, using 20% of the available chunks and at most 50,000 rows per selected chunk. Before scaling, each feature was clipped to its estimated mean ± 5 standard deviations; the saved centring, scaling and clipping values were reused for SAE inference.

**SAE architecture, initialization and optimization** Top-k SAEs were evaluated across the following dictionary sizes $D$ and sparsity levels $k$:

$$D \in \{2{,}302, 4{,}604, 11{,}520, 23{,}040\}$$

$$k \in \{16, 32, 64\}$$

For a preprocessed input vector x, the SAE first applied a linear encoder, rectified linear unit and hard top-k selection:

$$h = \mathrm{ReLU}(W_{\mathrm{enc}}^{\mathsf{T}} x + b_{\mathrm{enc}}), \qquad z = \mathrm{TopK}_k(h)$$

All but the k largest positive activations were set to zero. Reconstruction was then calculated as:

$$\hat{x} = W_{\mathrm{dec}}^{\mathsf{T}} z + b_{\mathrm{dec}}$$

Decoder rows were initialized from randomly selected, L2-normalized training vectors. Encoder weights were initialized as 0.1 $W_{\mathrm{dec}}^{\mathsf{T}}$, and encoder and decoder biases were initialized to zero.

Models were optimized with AdamW using an initial learning rate of $4 \times 10^{-4}$, $\beta_1$ = 0.9 and $\beta_2$ = 0.999. The configured batch size was 30,240. The learning rate was linearly warmed up over the first 10% of optimizer updates and then cosine-decayed to a minimum of $4 \times 10^{-5}$. Gradients were clipped to a global norm of 1.0.

The total objective was:

$$\mathcal{L} = \mathcal{L}_{\mathrm{Huber}}(x, \hat{x}; \delta = 5) + \frac{1}{1024}\mathcal{L}_{\mathrm{AuxK}} + 0.05(\bar{a} - 0.01)^2 + 0.0005\frac{1}{D}\sum_{j=1}^{D}(\|d_j\|_2 - 1)^2$$

Here, $\bar{a}$ is the fraction of active latent units in the batch and d_j is decoder dictionary vector j. The AuxK term reconstructed the main-reconstruction residual using 128 auxiliary features and the same Huber loss with $\delta = 5$. Decoder rows were explicitly renormalized every 200 optimizer updates. Feature inactivity was monitored using a 500-update threshold, and the implementation permitted dead-feature resampling after the warm-up period; the final selected model contained no dead features.

The operational D = 4,604, k = 64 checkpoint used to generate the current semantic-motif system was trained for 20 epochs and recorded 7,800 optimizer updates. Final-epoch values were 0.1105 for the main reconstruction loss, 0.0972 for the auxiliary reconstruction loss and 0.779 for explained variance, with a dead-feature fraction of zero. The configuration was carried forward based on joint inspection of reconstruction fidelity, feature utilization and the interpretability of the downstream token communities. These settings are summarized in Supplementary Table 2.

**FAISS token graph and Leiden community detection** The operational $D = 4{,}604$, $k = 64$ SAE projected residue representations from 109,220 proteins into 39,001,072 sparse token-activation vectors, which formed the vertices of the semantic-motif graph.

Before neighbour search, token-activation vectors were L2-normalized. Because the selected product-quantization layout required a compatible dimension, the 4,604-dimensional vectors were zero-padded to 4,608 dimensions. Approximate nearest-neighbour search used a FAISS inner-product IndexIVFPQ index; inner product is equivalent to cosine similarity after L2 normalization. The index used 32,768 inverted lists, 64 product-quantization subquantizers and 8 bits per sub-quantizer. Index training used a deterministic blockwise random sample of 2,000,000 token vectors with seed 42 and a sampling block size of 1,000,000. All 39,001,072 token vectors were added to the trained index in batches of 200,000 and searched in batches of 100,000 using `nprobe`=64, retrieving 50 neighbours per token.

Valid FAISS source-neighbour pairs were passed directly to an undirected weighted cuGraph graph, with cosine similarity as edge weight and a minimum edge-weight parameter of 0.0.

Weighted Leiden community detection used resolution 0.1 and the cuGraph defaults for iteration control and random state. It assigned the graph vertices to 364 communities with raw identifiers 0–363.

**Positional reindexing of the 364 Leiden communities** The raw Leiden identifiers were deterministically reordered to make motif identifiers reflect broad positional behavior along proteins. For these reindexing statistics, residue coordinates within a protein of length L were assigned using numpy.linspace(0, 1, L); proteins shorter than five residues were excluded from the positional-summary calculation.

For each raw motif, the mean and standard deviation of its relative residue positions were calculated together with its mean and maximum within-protein residue fractions. Motifs with positional standard deviation $\leq 0.2$ were categorized by mean relative position as follows:

- N-terminal: mean position $< 0.25$.
- Middle/conserved: mean position from 0.25 to 0.75, inclusive.
- C-terminal: mean position $> 0.75$.

Motifs with positional standard deviation $> 0.2$ were categorized as dominating when their mean within-protein fraction was $> 0.3$ or their maximum within-protein fraction was $> 0.8$; all remaining motifs were categorized as random/scattered. Motifs were ordered first by category - N-terminal, middle/conserved, C-terminal, dominating and random/scattered - and then by increasing mean relative position within each category. New semantic-motif identifiers 1-364 were assigned in this order. The resulting classes comprised 58 N-terminal, 151 middle/conserved, 128 C-terminal, 19 dominating and 8 random/scattered motifs.

The positional reindexing preserved all 364 Leiden communities in one-to-one correspondence.

**Residue assignment and motif-string construction** Each residue remaining after special-token removal corresponded to one graph vertex and inherited exactly one semantic-motif label from its Leiden community. The resulting protein representation was a complete residue-wise motif string.

For a protein of length L, its motif string was the ordered sequence:

$$M = (m_1, m_2, \ldots, m_L), \qquad m_i \in \{1, \ldots, 364\}$$

Consecutive repetitions of a motif label remained separate residue-level tokens.

**Motif n-gram profiles** Motif n-gram profiles were calculated at the AAI50-cluster level. Complete residue-wise motif strings of member proteins were concatenated in analysis-table order as one whitespace-delimited document per cluster. Adjacent protein strings therefore contributed junction-spanning bigrams and trigrams to the cluster profile.

Within each AAI50-cluster document, individual motif identifiers and adjacent pairs and triplets of identifiers were counted, including single-character identifiers. An n-gram was retained if it occurred in at least three AAI50 clusters in analysis groups containing three or more clusters, or in at least one cluster in smaller groups. The resulting counts were weighted by term frequency–inverse document frequency (TF-IDF), and each AAI50-cluster profile was normalized to unit L2 length.

##### Reference architecture signatures and open-set assignment

For reference anchoring, 13,504 reference major capsid proteins from NCBI Virus were processed using the same ESMCapsid-C layer-28 extraction, SAE projection, Leiden label system and motif-string procedure. Reference signatures were constructed from AAI50 clusters assigned to eight curated operational architecture categories: HK97-like, picorna-like, NCLDV-like, BTV-like, Microviridae-like, Geminiviridae-like, Circoviridae-like and levi-like.

Candidate AAI50 clusters were compared with each reference signature using weighted coverage of category-specific core n-grams together with global motif-profile similarity. A candidate was assigned to the highest-scoring category only when its score exceeded the corresponding category-specific open-set threshold. Cosine similarity and Jensen-Shannon divergence were retained as complementary confidence measures. Before Jensen-Shannon divergence was calculated, TF-IDF profile vectors were converted to unit-sum distributions. Thus, final category assignments integrated local enrichment of category-associated motif n-grams with global similarity of the complete motif profile.

##### Core, high-weight and driver motifs and positional profiling

Within each architecture category, motif prevalence was defined as the fraction of clusters in which a motif occurred. Core motifs had prevalence at least 0.95 and TF-IDF variance within the lower 40th percentile among high-prevalence motifs. High-weight motifs had mean TF-IDF at or above the 75th percentile within the category. Driver motifs were selected from high-weight motifs by ranking TF-IDF variance and retaining the smallest set explaining 50% of cumulative variance (driver50); an 80% cumulative-variance set was retained for sensitivity analysis. Shared drivers for a pair of clusters were identified from large values of the elementwise product of their TF-IDF vectors, whereas cluster-specific drivers were identified from the positive components of their vector difference.

Positional preferences of individual semantic-motif labels and motif n-grams were summarized separately from the positional reindexing procedure described in Supplementary Note (Additional details of sparse feature analyses). For an occurrence beginning at zero-based residue index i in a protein of length L, its relative position was defined as r = i/L. The interval [0,1) was divided into 20 equal-width bins. For an n-gram, i was the start index of the occurrence. Background profiles counted every residue position, whereas core and driver profiles counted occurrences of the corresponding motif labels or n-grams. Within each profile category, pooled bin counts were divided by the total number of occurrences in that category. No additional smoothing was applied.

##### Mapping conserved semantic motifs onto capsid shell structures

Conserved motif n-gram cores were mapped to three representative HK97-like assemblies and three representative picorna-like assemblies (Supplementary Table 3). The particle centre was defined as the mean coordinate of all C-alpha atoms in the assembly. For each residue, normalized radial position was calculated from its C-alpha distance to the particle centre and scaled between the assembly-wide minimum and maximum radii. Inner-, middle- and outer-shell positions were defined as normalized radial positions below 0.33, from 0.33 to below 0.67 and at least 0.67, respectively. Mapped core positions were intersected with structure-resolved residues in the representative chain and merged into contiguous ranges.

Observed radial spread was calculated as the standard deviation of normalized radial positions among mapped core residues. Spatial compactness was summarized by the median pairwise C$\alpha$ distance among mapped core residues with resolved coordinates. Local packing density was summarized as the mean number of resolved residues within 10 Å of each mapped core residue. In the final Source Data, these three statistics were compared with the corresponding distributions from 500 random residue sets for each displayed mapping. Empirical P values used the plus-one correction, $P = (b + 1)/(n + 1)$, where $b$ is the number of null statistics at least as extreme as the observed value and $n$ is the number of random sets.

Interface-related summaries compared mapped core residues with the remaining resolved residues in the same representative chain or entity. These summaries included the fraction of residues within 8 Å of another chain, the minimum C$\alpha$ distance to another chain and the number of residues from other chains within 12 Å. Recurrent protomer geometry was assessed after projecting representative-chain C$\alpha$ coordinates into a tangent-plane coordinate system. The principal axis of the mapped core residues defined the band direction, and the perpendicular axis defined the cross-band direction. Band width was quantified as the standard deviation along the cross-band axis, whereas anisotropy was calculated as the ratio of the standard deviations along the band and cross-band axes. These statistics were compared with 100 null residue sets that preserved residue number and contiguous segment lengths and matched segment-level radial positions using tolerance values of 0.03, 0.05, 0.08, 0.12 and 0.20 and random seed 7.

### Supplementary Table 1

#### Supplementary Table 1 | Protein language models benchmarked in this study.

| Model | Checkpoint / version | Developer | Model size | Model source used | Training data / modalities | Output used in this study |
|---|---|---|---|---|---|---|
| ESM2-650M | facebook/esm2_t33_650M_UR50D | Meta AI / FAIR | 650M; 33 layers | Synthyra/ESM2-650M | Protein sequences from UniProt 2021_04 | Residue- and sequence-level embeddings |
| ESM3-open | esm3-sm-open-v1 / esm3-open | EvolutionaryScale / Biohub | 1.4B | biohub/esm3-sm-open-v1 | Sequence, structure and function tracks; natural-protein, structure and function-annotation corpora | Sequence representations |
| ESMC-600M | esmc-600m-2024-12 | EvolutionaryScale / Biohub | 600M; 36 layers | Synthyra/ESMplusplus_large | Protein sequences from UniRef, MGnify and JGI, clustered at 70% identity | Sequence embeddings and base checkpoint for ESMCapsid models |
| ProFluent-E1-600M | Profluent-Bio/E1-600m | Profluent | 600M | Synthyra/Profluent-E1-600M | Protein sequences from the Profluent Protein Atlas, with optional homologous context | Sequence embeddings |

### Supplementary Table 2

#### Supplementary Table 2 | Key ESMCapsid and sparse-autoencoder settings.

| **Component** | **Input / base model** | **Objective / architecture** | **Training length / selection** | **Key settings** | **Role** |
|---|---|---|---|---|---|
| ESMCapsid-S adaptation | ESMC | Supervised contrastive LoRA | 10 epochs | AdamW; learning rate $1 \times 10^{-5}$; weight decay 0.01; batch size 32; BF16 | Remote capsid detection |
| Frozen probe heads | Mean-pooled frozen embeddings | Two-layer MLP; hidden widths 128 and 256; ReLU; dropout 0.1 | Task-specific | Class weighting used when label imbalance was substantial | Binary detection and biological annotation probes |
| Layer-16 two-stage screen | ESMCapsid-S layer 16 | Normal head followed by hard-negative head | Fixed validation thresholds | Normal-head threshold 0.99; hard-negative-head threshold 0.9675 | Metagenomic-scale screening |
| ESMCapsid-C continued pretraining | ESMC | Full-parameter masked-language modelling; 15% masking | 100,000 steps | AdamW; learning rate $1 \times 10^{-5}$; >210,000 capsid proteins | Capsid-specific representation learning |
| Operational sparse autoencoder | Layer-28 token representations (1,152 dimensions) | Top-k SAE; D = 4,604; k = 64; Huber, AuxK, activity-rate and decoder-norm terms | 20 epochs; 7,800 optimizer updates | AdamW; learning rate $4 \times 10^{-4}$ to $4 \times 10^{-5}$; batch size 30,240; 4,000,000 sequence-balanced draws per epoch | Token-level sparse activations and semantic-motif construction |

### Supplementary Table 3

**Supplementary Table 3 | Representative capsid assemblies used for motif-to-structure mapping.**

| Architecture category | PDB ID | Representative chain(s) | Use |
|---|---|---|---|
| HK97-like | 1OHG | A | Experimental assembly |
| HK97-like | 2XYY | A | Experimental assembly |
| HK97-like | 5UU5 | G | Experimental assembly |
| picorna-like | 1B35 | A, B and C | Three capsid-protein chains |
| picorna-like | 1HXS | 1, 2 and 3 | Three capsid-protein chains |
| picorna-like | 5WTE | A, B and C | Three capsid-protein chains |
